# A nicotine biosensor derived from microbial screening

**DOI:** 10.64898/2026.03.11.710934

**Authors:** Uroš Kuzmanović, Mingfu Chen, Roger Charles, Abdurrahman Addokhi, Margarita A. Tararina, Kristin A. Hughes, Anthony M. DeMaria, Prerana Sensharma, Anant Gupta, Sreyash Dasari, Nelson Leonardo Gonzalez Dantas, Karthika Sankar, Zhehao Zhang, Hao Zang, Karen N. Allen, Catherine M. Klapperich, Mark W. Grinstaff, James E. Galagan

**Author notes:** These authors contributed equally (Uroš Kuzmanović, Mingfu Chen).

## Abstract

Physiologically relevant biosensors are in increasingly high demand, yet existing ones are severely limited in the number and type of biomarkers that are detected. The lack of biorecognition elements for most medically relevant biomarkers restricts the development of next generation single and continuous use monitors. Over billions of years, microbes have evolved a vast array of proteins to sense and metabolize small molecules, including those pertinent to human health. Of particular interest to us is the identification and subsequent integration of new microbial redox enzymes into electronic biosensors building off the established electrochemical technology of the continuous glucose monitor. Here we deploy genomic screening to identify analyte specific redox enzymes for biosensor development. As a proof of concept, we report the first electrochemical enzyme-based nicotine biosensor from a novel microbial enzyme, and use a variant with improved catalytic performance to enhance sensor performance. The biosensor detects nicotine over 0.4-100 μM, a range relevant to nicotine concentrations present in active smoker sweat, saliva, gastric juice, and urine. This microbial mining approach for discovering redox enzymes expands the sensing parts toolbox available over conventional antibodies and aptamers.

## Introduction

Single and continuous use biosensors are improving patient care by enabling quantitative measurements to inform medical decision-making. Continuous use biosensors will play an even greater role in healthcare as we advance towards real-time monitoring of physiology^2–10^. However, we are only able to detect a small fraction of the physiologically relevant biomarkers. This limitation is a result of the lack of biorecognition elements for biosensor development. The majority of current biorecognition elements comprise antibodies and aptamers, which are ideal for single use applications but are fraught with difficulties for use in continuous measurements. The benchmark for continuous biosensing is the continuous glucose monitor (CGM) which utilizes a redox enzyme, glucose oxidase, as the biorecognition element^11–14^. Glucose oxidase (GO_X_) catalytically degrades glucose and liberates electrons^13,15^, thus transducing a biological signal into an electrical signal^13,16,17^. While enzyme-based electrochemical sensors are known for a small group of analytes including lactate, cholesterol, uric acid, urea, and acetylcholine^18–22^, sensors for other physiologically and medically relevant analytes are lacking, primarily due to the lack of analyte specific redox enzymes. Consequently, there is a need to develop generalized methodologies to identify new enzymes for continuous use biosensors.

Microorganisms are an enormous and untapped reservoir for identifying molecular sensing parts for biosensors. Microbes have evolved over 3 billion years to detect and respond to virtually all stimuli relevant to human biology, biotechnology, and our environment^23^. Microbial proteins, like glucose oxidases are particularly attractive for biosensing as they are naturally amenable to bacterial or cell free expression systems and, thus, easier and cheaper to produce. Eukaryotic proteins, by contrast, often require post-translational modifications, complex chaperoned folding, and multimeric assembly that is more difficult to replicate. Similarly, bacterial proteins are more amenable to optimization for biosensing through protein engineering and directed evolution. In prior work we demonstrated the use of genomic screening to identify analyte specific allosteric transcription factors (aTFs)^24^ and developed a novel class of biosensors based on aTFs^24–31^. Here we extend the screening of microbes to identify analyte-specific redox enzymes.

As a proof of concept, we report here a novel real-time continuous biosensor for nicotine. Nicotine is the leading cause of preventable deaths worldwide and the addictiveness of nicotine is the primary reason for tobacco abuse^32,33^. Nicotine monitoring is important for applications including detection of nicotine exposure from emerging nicotine delivery devices (e.g., nicotine pouches, vaping, and e-cigarettes)^34^, protection of vulnerable populations such as children and pregnant women from second-hand smoking^32^, and cessation of smoking^35^. The gold standard for nicotine analysis is mass spectrometry (GC/LC-MS)^36,37^ which has high specificity, sensitivity, and analytical accuracy but requires complex instrumentation, trained personnel, and long turnaround times. While alternative nicotine biosensors exist^38–49^, these biosensors lack specificity due to the use of sensing parts that do not selectively target nicotine. Herein, we report a highly specific nicotine electrochemical biosensor with micromolar sensitivity developed from a microbial redox enzyme, nicotine oxidoreductase (NicA2) recovered from our screening of *Pseudomonas putida* – a microbe present in the soil in a tobacco field^50,51^. Further, we demonstrate enhanced sensor performance using a protein variant with improved catalytic activity. The resulting biosensor detects nicotine over a range of 0.4-100 μM relevant to nicotine concentrations present in active smoker sweat, saliva, gastric juice, and urine. The biosensor also performs real time, continuous monitoring of nicotine from artificial sweat.

## Results

### Identification Of NicA2 Using a Microbial Genomic Screen

We screened the bacterium *Pseudomonas putida S16* (*P. putida S16*), known to have the ability to metabolize nicotine^52^ by adapting our previously successful genomic screening strategy^24^. Building on the knowledge that metabolic genes, often encoding analyte-specific enzymes, are found in genomic clusters we prioritized their identification in this application. Using our approach, we identified a gene cluster that was significantly up-regulated in the presence of nicotine (Figure 1A-B) that we termed the nicotine responsive genome island (NRGI). The NRGI contains a total of 48 genes and overlaps the previously reported gene cluster associated with nicotine catabolism in *P. putida S16*^53^ (Figure 1C). The NRGI includes several genes of interest for biosensor development including: 1) a predicted but uncharacterized AraC family transcription factor (TF) that was the only significantly upregulated TF (>3-fold change in expression) in the presence of nicotine during lag-phase of cell culture and 2) the 2^nd^ gene in an upregulated cluster which encodes a nicotine oxidoreductase (NicA2) (16-fold change) (Figure 1D).

**Figure 1.**
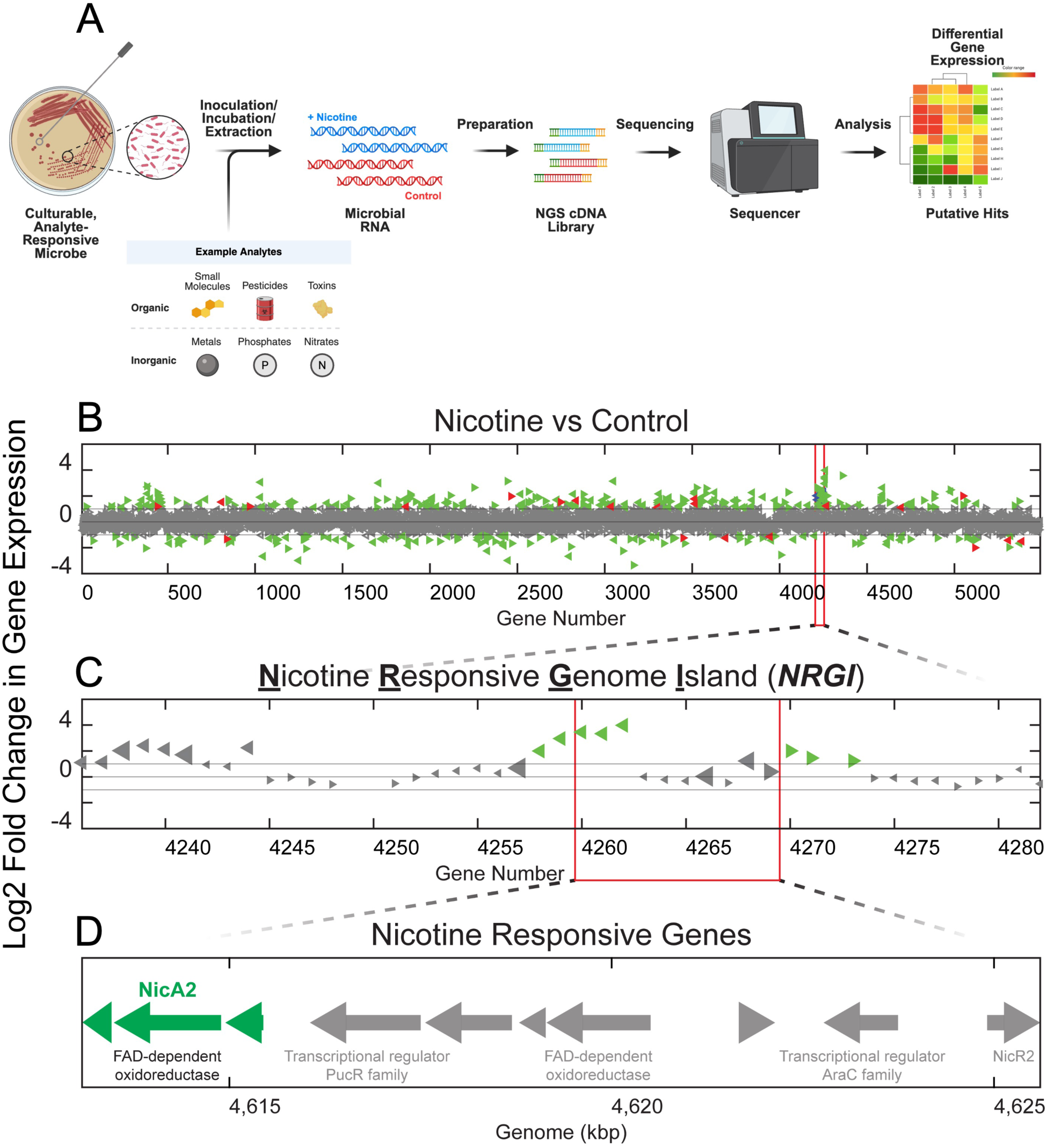
Identification of nicotine responsive genes from *P. putida S16* using genomic screening. **(A)** Schematic of our genomic screening pipeline utilized to identify a nicotine degrading gene cluster from ***P. putida S16***. **(B)** Genome-wide log2-fold change gene expression in response to nicotine relative to control during lag-phase microbial growth. Each triangle is a gene (red = differentially expressed TF, blue = gene with annotated analyte related function, green = differentially expressed other gene, gray = non-differentially expressed other gene. Horizontal lines above and below 0 indicate +/- 1 log2 fold change expression). **(C)** Zoomed-in view of the Nicotine Responsive Genome Island (NRGI). **(D)** Nicotine responsive genes include *nicA2*, which was the highest differentially expressing redox enzyme.

NicA2 has been characterized as a FAD-dependent enzyme (MW ∼53 kDa) that catalyzes the first step of nicotine degradation in *P. putida S16*^54,55^. Biochemical characterization has previously demonstrated that NicA2 is extremely specific to nicotine (K_M_ ∼100 nM), highly stable, and can use oxygen as electron acceptor^55,56^. These properties suggest the potential for using NicA2 as the biorecognition element for electrochemical biosensing analogous to GO_X_-based biosensing. Although NicA2 has been studied as a potential therapeutic for smoking cessation^56,57^, the enzyme has not been previously utilized as a biosensor and little is known about its electrochemical behavior.

### NicA2 Biochemical Characterization *In Vitro*

We isolated the NicA2 protein and biochemically confirmed its activity. The gene encoding NicA2 was cloned into a recombinant vector with histidine tags and heterologously expressed in *E. coli BL21.* The protein was then purified using fast protein liquid chromatography (FPLC) via size-exclusion chromatography (SEC). NicA2 activity was verified using the Amplex UltraRed (AUR) assay (Figure 2A). NicA2 catalyzes the oxidation of nicotine into N-methylmyosmine in a FAD-dependent reaction^55,58^. In the AUR assay, the reduced NicA2 is regenerated by O_2_ and produces H_2_O_2_, which is converted by a peroxidase to the fluorescent reporter resorufin. By varying nicotine (Figure 2B) and enzyme concentrations (Figure 2C), we showed that the amount of H_2_O_2_ production is both substrate- and enzyme-dependent. We characterized the stoichiometry of the redox reaction catalyzed by NicA2 showing a H_2_O_2_: nicotine ratio of 1:1 (Figure 2D), confirming that 2 electrons are transferred from nicotine during each reaction cycle. H_2_O_2_ production from nicotine required NicA2, and consistent with this, H_2_O_2_ production was significantly reduced in the presence of myosmine, a product analog and NicA2^55^ inhibitor (Figure 2E).

**Figure 2.**
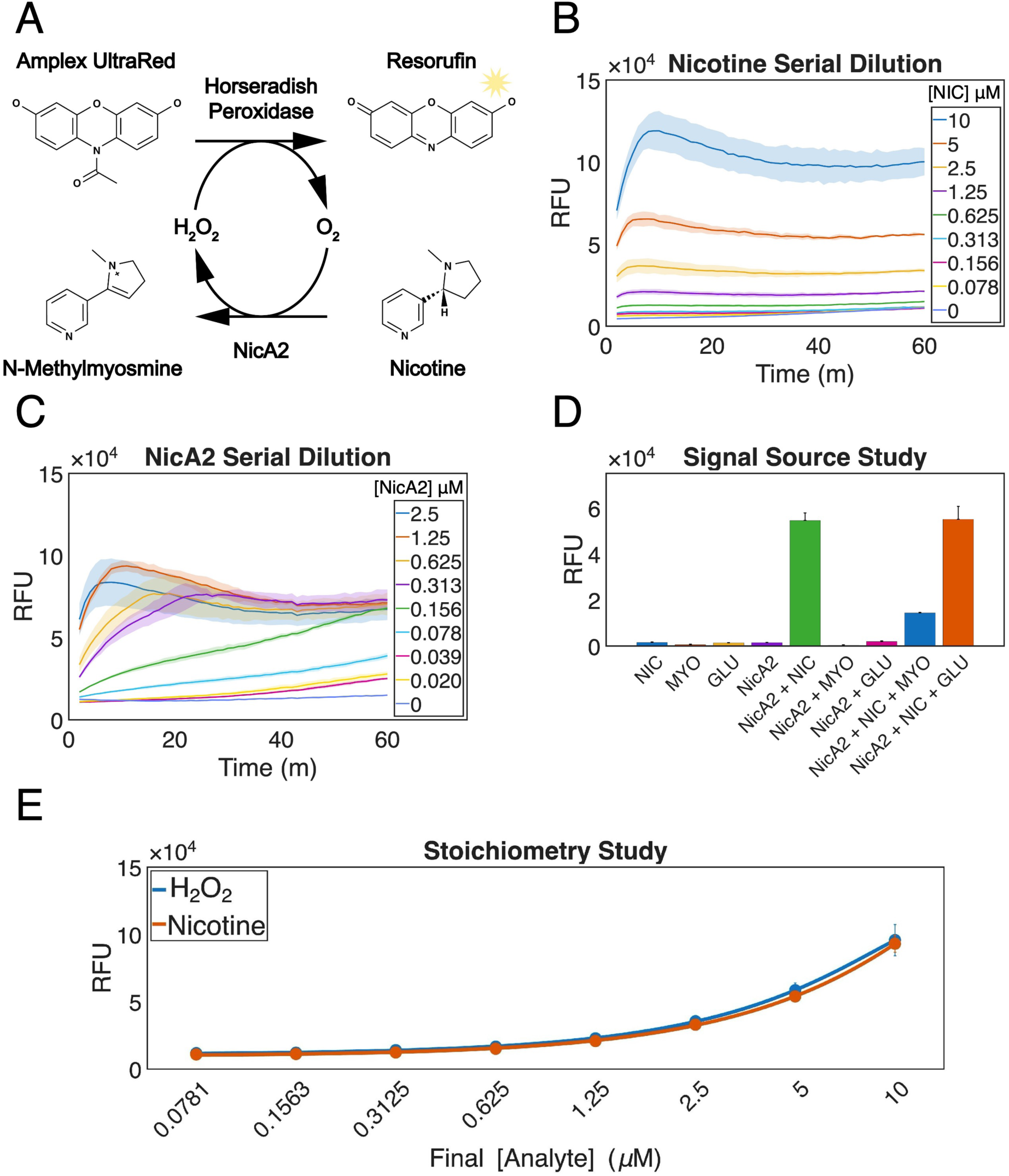
Biochemical characterization of NicA2. **(A)** Amplex UltraRed assay for characterizing nicotine oxidation by NicA2. **(B)** Nicotine oxidation is analyte dependent at constant enzyme concentration. **(C)** Nicotine oxidation is enzyme concentration dependent. **(D)** Comparison of nicotine and H_2_O_2_ assay results at equimolar concentrations confirms a 1:1 production of H_2_O_2_ from nicotine by NicA2. **(E)** H_2_O_2_ production requires nicotine and NicA2. Nicotine is not independently oxidized without NicA2. NIC: nicotine; MYO: myosmine; GLU: glucose.

### Engineering and Optimizing a NicA2 Electrochemical Sensor

The stoichiometric production of H_2_O_2_ by NicA2 provided the opportunity to develop a sensor analogous to GO_X_-based glucose sensors^59^. For GO_X_, oxidation of glucose transfers two electrons to reduce an enzyme-bound flavin adenine dinucleotide (FAD) cofactor. Additional rounds of catalysis require the regeneration of oxidized FAD. In first-generation electrochemical sensors, electrons are transferred from FAD to molecular oxygen (O_2_) to produce hydrogen peroxide (H_2_O_2_). Prussian Blue (PB) then acts at the electrode surface to catalyze the reduction of H_2_O_2_, and oxidized PB is subsequently reduced at the electrode surface, thus generating an electrode current^60,61^. We developed a first-generation electrochemical nicotine biosensor using this same principle (Figure 3A). As NicA2 was not previously studied for biosensing, we systematically tested multiple sensor parameters to optimize sensor performance, including the enzyme immobilization, screen-printed electrode design, and enzyme amount.

**Figure 3.**
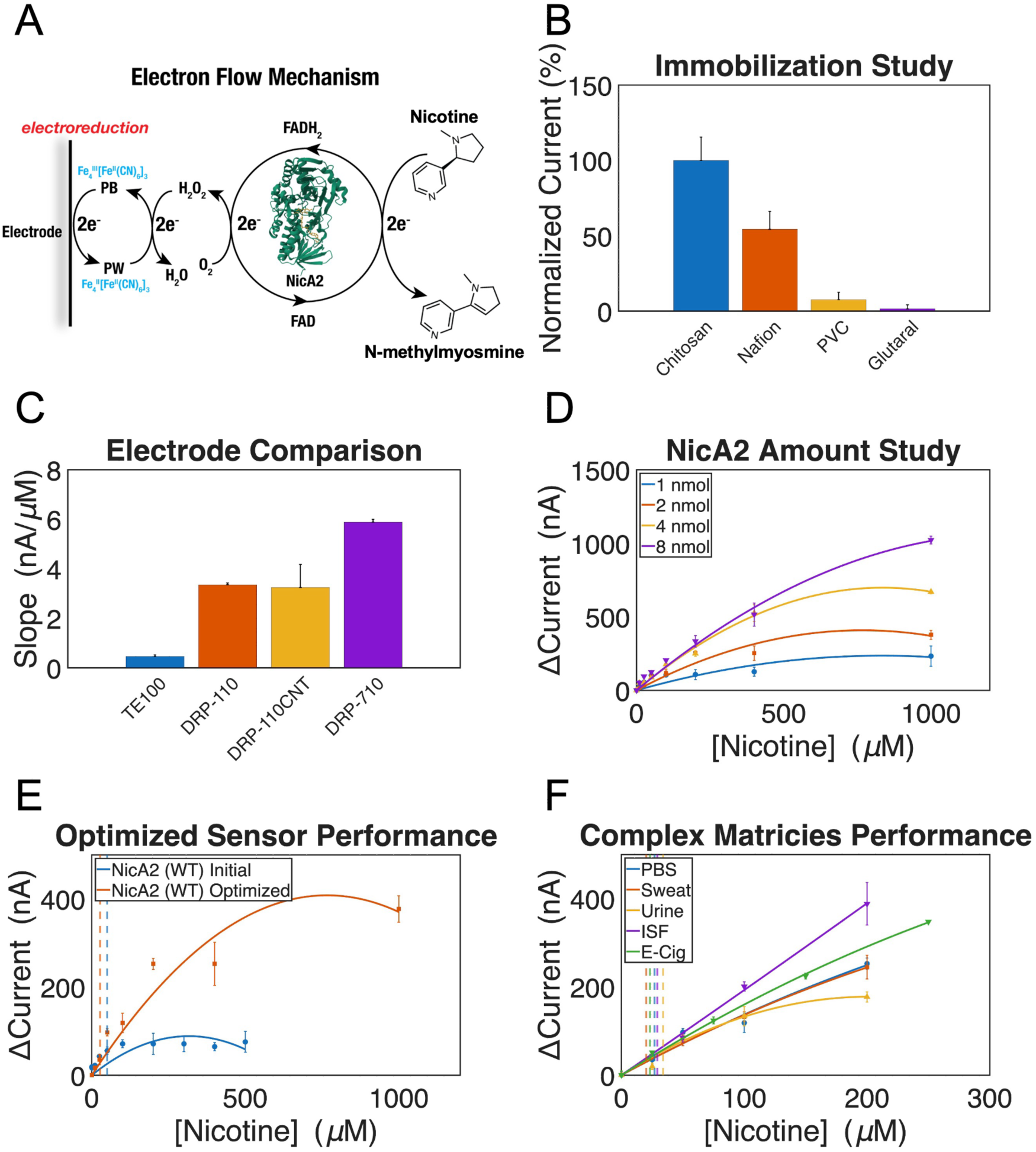
Engineering an electrochemical nicotine biosensor using WT NicA2. **(A)** Electron flow mechanism for a first-generation electrochemical NicA2 biosensor. FAD: flavin cofactor; (PB) Prussian Blue, (PW) Prussian White. **(B)** Comparison of immobilization methods of enzyme. Currents were measured with 400 μM nicotine in PBS added to sensor prepared with different immobilization methods. Currents were normalized against 1 wt% chitosan in 0.5 wt% acetic acid. **(C)** Comparison of screen-printed electrode designs. TE100 with medium PB, DRP-110 with thick PB, DRP-110CNT with thick PB, and DRP-710. **(D)** Optimization of enzyme deposition amount. Dose-response curves for nicotine detection in PBS with varying enzyme quantity. **(E)** Comparison of optimized sensor and initial sensor. Initial sensor: TE100 electrode, 1 nmol NicA2, immobilization with 1 wt% chitosan in 0.5 wt% acetic acid. Optimized sensor: DRP-710 electrode, 2 nmol NicA2, was immobilization with1 wt% chitosan in 0.5 wt% acetic acid. Dashed lines indicate the limit of detection (LOD) defined as the nicotine concentration with a signal greater than 3x pool standard deviations above background. Initial and optimized sensor LOD values are 49.9 μM and 26.5 μM, respectively. **(F)** Dose-response curves of optimized sensor for nicotine detection in PBS, artificial interstitial fluid, artificial sweat, human urine, and e-cigarette liquid. Dashed lines indicate the limit of detection. LODs in PBS, artificial interstitial fluid, artificial sweat, human urine, and e-cigarette liquid are 26.5 μM, 29.3 μM, 20.2 μM, 34.2 μM, 23.2 μM, respectively.

First, we compared four different strategies for immobilizing NicA2 on an electrode surface. All testing was performed using screen-printed electrodes (SPEs) deposited with Prussian Blue (PB) (TE100 electrodes commercially acquired from CH Instruments, Inc). Initially, we tested chitosan dissolved in different acetic acid solutions. We selected chitosan for its membrane-forming ability, water permeability, adhesion capability, hydrophilicity, and biocompatibility^62–64^. We monitored the responses to nicotine by measuring the current elicited upon addition of 400 μM nicotine solution. Use of a lower acetic acid weight percent (0.5 wt%) improved device performance likely due to higher enzymatic activity (SI Figure 20A)^55^. We then tested three additional immobilization strategies: 1) Nafion as a protective layer to exclude potential interferents away from the transducer surface^65,66^; 2) bovine serum albumin covered with polyvinyl chloride (PVC)^67–69^ to stabilize the enzyme by hydrophobic interactions; and 3) glutaraldehyde as a cross-linking reagent for the enzyme layer^70–74^. Although all three methods generated an output signal in the presence of nicotine, the biosensor prepared with 1 wt% chitosan in 0.5 wt% acetic acid yielded the highest current response. (Figure 3B and SI Figure 20B). Thus, we adopted the immobilization method employing NicA2 and 1 wt% chitosan in 0.5 wt% acetic acid for all subsequent testing.

Second, we assessed four different commercially acquired SPEs: TE100, DRP-110, DRP-110CNT, and DRP-710. TE100 possesses a carbon working electrode (diameter=3 mm), a carbon counter electrode, and an Ag/AgCl pellet reference electrode. Compared to TE100’s smaller active surface, DRP-110, DRP-110CNT, and DRP-710 possess larger active surface (diameter=4 mm). DRP-110, DRP-110CNT, and DRP-710 possess working electrode made of carbon, multi-walled carbon nanotubes, and Prussian blue/carbon, respectively. For TE100, DRP-110, and DRP-110CNT SPEs, we electrodeposited different thicknesses of PB on electrodes. DRP-710 SPEs were used without further PB deposition. We compared the performance of each electrode by comparing the current elicited after addition of H_2_O_2_. Across all the electrodes and PB thicknesses, current positively correlated with H_2_O_2_ concentration within the tested range of 0 to 800 μM (SI Figure 20). Carbon SPEs (TE100, DRP-110, and DRP-110CNT) displayed statistically significant PB thickness-dependent increases in the slope of this response (Figure *3*C), suggesting that more active Prussian Blue sites are accessible to hydrogen peroxide with increasing layers of Prussian Blue on the carbon surface^75,76^. However, DRP-710 exhibited the highest sensitivity and current density across the four SPEs tested (SI Figure 20, SI Figure 20I). Thus, we selected DRP-710 for the subsequent NicA2 sensor development.

Third, we evaluated the impact of using different amounts of NicA2. Using chitosan immobilization and DRP-710 SPEs, we conducted chronoamperometric measurements with 1, 2, 4, and 8 nmol of deposited NicA2. Across all enzyme amounts, current positively correlated with nicotine concentrations between 0 and 1000 μM (Figure 3D). At 1000 μM, the current from sensors with 1 and 2 nmol NicA2 did not reach baseline within 2 hours indicating incomplete depletion of analyte (SI Figure 22A-B). In contrast, 4 and 8 nmol did result in apparent analyte depletion (SI Figure 22C-D). To gauge the impact of NicA2 concentration upon sensitivity, we performed linear regression analysis on the dose-response curves in the range of 0 to 200 μM, as this range approximates the physiological level of nicotine^77^. Relative to 1 nmol, statistically significant increases in sensitivity was observed for 2 and 4 nmol, and 8 nmol. However, the largest increase in sensitivity per nmol of enzyme was observed between 1 and 2 nmol (SI Table 6). To balance sensitivity against the use of more enzyme, we selected 2 nmol enzyme loading for subsequent sensor development.

### Characterization of the Optimized NicA2 Electrochemical Sensor in Complex Matrices

Based on the above results, we selected the following as the configuration for our final WT NicA2 biosensor: immobilization with 1 wt% chitosan in 0.5 wt% acetic acid; DRP-710 electrode; 2 nmol deposited NicA2. To assess the impact of optimized configuration, we compared electrode performance to an initial sensor using TE100 electrodes and 1 nmol of NicA2 immobilized with 1 wt% chitosan in 0.5 wt% acetic acid. The optimized sensor displayed a ∼4-fold increase in signal (252.66 nA vs. 64.66 nA for optimized vs. initial) and ∼2-fold decrease in LOD (26.5 μM vs. 49.9 μM) (Figure 3E). We then characterized the LOD of the optimized sensor in different complex matrices. Testing in bodily fluids resulted in LODs of 26.5 μM, 29.3 μM, 20.2 μM, 34.2 μM, in PBS, artificial interstitial fluid, artificial sweat, and human urine respectively, while testing in e-cigarette liquid resulted in an LOD 23.2 µM (Figure 3F).

The performance of the sensor in e-cigarette liquid suggests the potential for use in monitoring nicotine delivery in e-cigarettes. To validate this use case, we first confirmed that sensor performance was not impacted by the presence of common interferants in e-cigarette liquid (SI Figure 25). We then tested accuracy of nicotine measurement on e-cigarette liquid from Juul^®^ pod diluted to different nicotine concentrations (SI Table 5). The recovery rate was 101.9, 104.3, and 99.4 % for concentrations of 50, 100, 200 μM, respectively. The corresponding relative standard deviation (RSD) was 6.6, 6.2, and 4.2 %. In addition to prediction accuracy, our sensor is also significantly less costly than existing methods for nicotine measurement such as mass spectrometry (SI Table 8).

### Engineering a Kinetically Improved Nicotine Biosensor with Mutated NicA2 [N462H]

Although applicable to nicotine delivery monitoring, the optimized nicotine sensor did not cover the range of nicotine in smoker sweat and urine (SI Table 3). To improve the sensitivity of our nicotine biosensor further, we utilized a previously reported rationally designed NicA2 enzyme variant^1,55,58,78^. As previously reported, a sequence similarity network was calculated, grouping NicA2 homologs^79^ into isofunctional clusters. One such cluster included monoamine oxidases (MAOs) with high catalytic efficiency^78^. Using multiple sequence alignments mapped to experimentally determined protein structures, N462 flanking the isoalloxazine ring, was identified as a position that differed between NicA2 and the monoamine oxidase homologs, with the efficient oxidases having hydrophobic/aromatic residues. A number of rationally designed protein variants were made at that position with the goal of improving oxygen reactivity. One variant in particular, N462H, was shown by transient-state kinetics to have an oxidation rate *k*_ox_ = 17 M^−1^s^−1^ compared to 2.2 M^−1^s^−1^ for WT NicA2, a ∼10-fold kinetic improvement. We compared this variant to WT NicA2 biochemically and confirmed a faster rate of nicotine conversion (Figure 4A).

**Figure 4.**
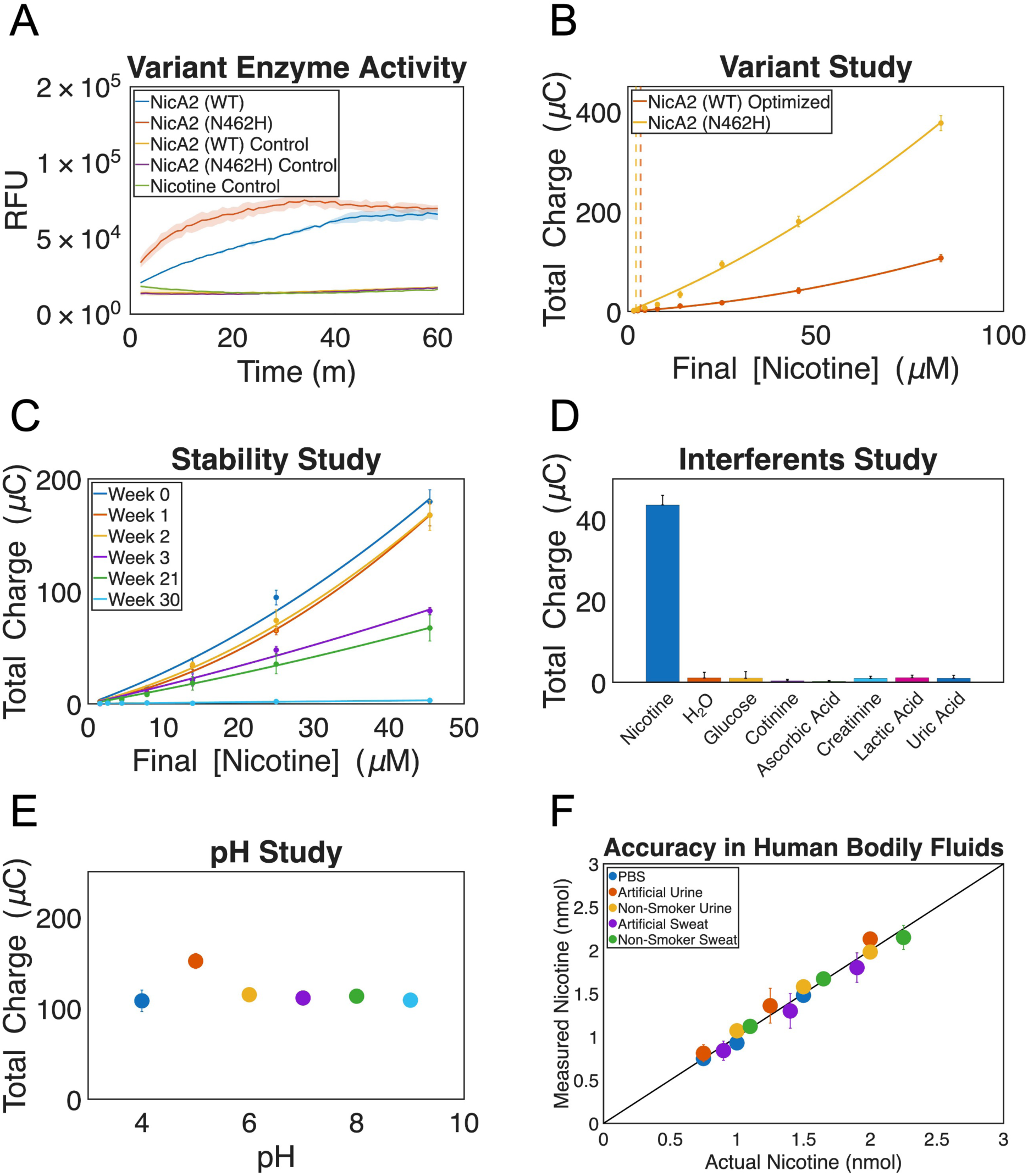
Characterization of an optimized electrochemical nicotine biosensor using a rationally designed NicA2 variant (N462H). **(A)** NicA2 (N462H) variant produced H_2_O_2_ at a faster rate than WT NicA2 at equimolar nicotine and enzyme concentrations. **(B)** Increased current output at equimolar nicotine concentrations using variant N462H NicA2, as compared to WT NicA2. **(C)** A stability study demonstrated biosensor stability in current output up to two weeks of storage at room temperature. **(D)** The nicotine biosensor showed little cross-reactivity to common interferents present in urine and sweat. **(E)** A pH study from 4-9 demonstrated that the nicotine biosensor is operational across this physiological range. **(F)** The nicotine biosensor was able to measure blindly doped artificial and human urine and sweat samples accurately.

Based on the hypothesis that increased enzymatic catalytic rate would translate to enhanced sensor performance, we engineered and characterized a sensor using N462H with the same configuration as that of wild-type NicA2 sensor. We compared the sensitivity of the N462H biosensor against WT NicA2 on physiologically relevant levels of nicotine spanning 1.56 – 83.33 μM (Figure 4B). The analysis revealed a modest improvement in LOD (2.03 µM and 2.14 µM for N462H vs WT) but a significant improvement in the limit of quantification (LOQ) (3.36 µM vs. 10.58 µM for N462H vs. WT). We found the nicotine biosensor to be stable for up to 2 weeks of storage at room temperature, showing only a slight decline in signal output (Figure 4C). There was a noticeable decrease in signal by the third week. Surprisingly, 21 weeks of storage produced a dose response similar to that of 3 weeks of storage while 30 weeks resulted in no signal output. Given the intended clinical and personal use of monitoring nicotine from human bodily fluids, we next determined the performance of the nicotine biosensor in the presence of common interferants in urine and sweat including the most common nicotine metabolite in humans, cotinine (Figure 4D)^80^. These interferents did not produce a statistically significant signal. A pH study from pH 4 – 9 using equimolar N462H NicA2 and 50 µM final nicotine showed that the biosensor was functional and uniform across all pH values with slightly higher output at pH 5 (Figure 4E).

Finally, the accuracy of the biosensor in urine and sweat was assessed by measuring blindly doped samples (Figure 4F). The biosensor was calibrated with three known concentrations of nicotine followed by the undoped complex fluid alone and then three blindly doped fluids. The mean accuracy was 100.54% with a mean standard deviation of 5.69% with no measured sample deviating more than 9.14% from the actual concentration.

### Continuous and Real-Time Monitoring

An advantage of enzyme based electrochemical sensors, as compared to affinity-based sensors using antibodies or aptamers, is the intrinsic ability for multiple and continuous measurement without the need for sensor regeneration. To demonstrate continuous real-time nicotine measurements, we deployed our sensor in a device that we designed called the WearStat (Figure 5A-B). The WearStat is a potentiostat built with a 3D printed ABS plastic housing, a coin-sized chargeable battery, and an Arduino circuit board recording current levels and relaying information wirelessly onto a custom-built MATLAB GUI. A flexible PDMS layer houses a paper channel which wicks sweat over the biosensor and into a large sink. Our results (SI Figure 18 and SI Figure 19), as well as previously reported ones^81^, demonstrate that increasing flow rate results in a higher current response. We thus designed the paper channel dimensions and material to maximize flow rate while decreasing diffusion layer thickness. For nicotine sensing we compared N462H deposited on an SPE with larger working electrodes (DS710BIG) and observed markedly improved current output with the same nicotine concentration over WT NicA2 with SPEs with smaller working electrodes (DS710). The LOD and LOQ of the N462H DS710BIG biosensor were found to be 0.37 µM and 3.69 µM respectively (Figure 5C). We then validated continuous nicotine sensing with this sensor deployed in WearStat in three different conditions. First, we measured nicotine concentrations from 50 – 1000 µM (0, 50, 200, 1000, 200, 50, 0 µM) in PBS from sponges in a continuous and stepwise fashion (Figure 5D). Second, physiologically relevant nicotine concentrations of 2 – 50 µM (0, 2, 10, 50, 10, 2, 0 µM) in PBS were measured using the same experimental approach (Figure 5E). Then, nicotine of concentrations 2 – 50 µM (0, 2, 10, 25, 50 µM) in both PBS and artificial sweat were measured sequentially over four cycles for more than six consecutive hours each (Figure 5F). Finally, to demonstrate we could continuously measure nicotine in human sweat, we measured nicotine of concentrations 2 – 50 µM (0, 2, 10, 25, 50 µM) in both PBS and doped non-smoker sweat sequentially over three cycles (Figure 5G). Although we saw a notable signal reduction in human sweat as compared to PBS, we observed reproducible signal responses over multiple cycles of step changes in nicotine concentration.

**Figure 5.**
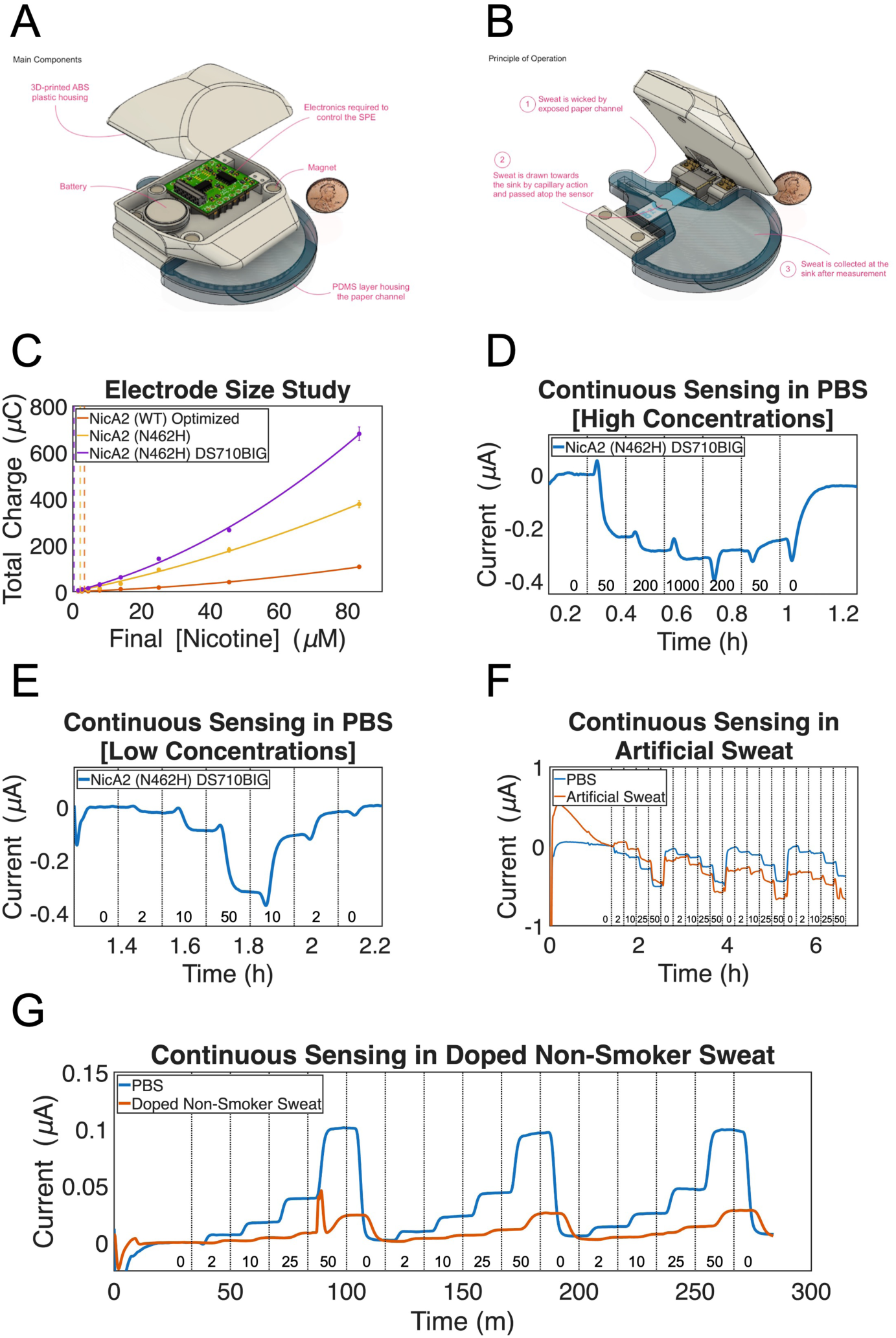
A wireless potentiostat, the WearStat, was developed to monitor nicotine levels quantitatively and in real-time from human sweat. **(A)** The WearStat device uncovered showing the electronic circuitry and battery next to a penny for scale. **(B)** The principle of operation for the WearStat device. 1) If adhered to a participant with sweat on their skin, the sweat will be wicked when in contact with the end of the paper channel. 2) Sweat is drawn to the sink by capillary action and passed above the biosensor. 3) Sweat is collected in the sink which is discarded or replaced after measurement. **(C)** Larger working electrodes (DS710BIG), and NicA2 N462H, drastically improve the current output with the same nicotine concentration over smaller working electrode SPEs (DS710) with WT NicA2. **(D)** The WearStat was used to measure nicotine concentrations in PBS from 50-1000 µM continuously and in real-time. **E)** The WearStat was used to successfully measure nicotine concentrations in PBS from 2-50 µM continuously and in real-time. **(F)** A four-cycle step function from 2-50 µM was measured in both PBS and artificial sweat. **(G)** A three-cycle step function from 2-50 µM was measured in both PBS and doped non-smoker sweat.

## Discussion

In this manuscript, we report the development and deployment of a highly specific and sensitive continuous biosensor for nicotine. We developed the biosensor from a microbial redox enzyme, NicA2, identified and recovered using a genomic screen applied to the nicotine degrading bacteria *P. putida S16*. The biosensor is the first enzyme-based electrochemical sensor for nicotine. We verified the ability of NicA2 to operate in an electrochemical sensor context and systematically tested different sensor parameters to optimize sensor performance. We further demonstrated the potential for improving sensor performance through the use of a rationally designed variant of NicA2 with enhanced catalytic activity. We validated the accuracy of our sensor in an array of benchtop tests including a test for accuracy in two use cases: monitoring nicotine delivery in e-cigarette liquid and measuring nicotine in volunteer non-smoker doped urine and sweat. We also demonstrate real-time continuous measurement of nicotine using a portable wearable potentiostat.

Our sensor exploits the capability of microbes that sense and degrade nicotine. The microbial metabolism of nicotine occurs by several distinct pathways in different bacterial species and includes the pyridine pathway found in gram-positive bacteria (*Arthrobacter nicotinovorans*)^51^, the pyrrolidine pathway found in gram-negative bacteria (*Pseudomonas putida*)^53^, and a variant of the pyridine and pyrrolidine pathway, found in *Ochrobactrum* and *Agrobacterium* species^82,83^. All three pathways depend on enzyme-mediated redox reactions to degrade nicotine. Although the proteins involved in this metabolism have been studied for roles in bioremediation and nicotine detoxification^56,84^, they have not previously been used for biosensor development.

More generally, our results highlight the potential of microbes as reservoirs for biorecognition elements for novel biosensors that can eventually be deployed on-body for real-time sensing. In prior work, we leveraged the sensing capabilities of microbes to develop a novel progesterone sensor based on a microbial transcription factor^24,26,27^. Here we extend this approach to the identification and harvesting of analyte-responsive redox enzymes. With advances in metagenomic screening^85,86^ and AI-powered *in silico* screening and protein engineering^87^, mining the functional diversity of microbes provides a solution to the need for more diverse biosensors for wearable health devices.

## Materials and Methods

### Measurements

Microplate experiments were analyzed by Spectramax M5 microplate reader (Molecular Devices, LLC, San Jose, CA). Electrochemical experiments were performed with VersaSTAT MC (Ametek Inc., Pennsylvania, PA). Scanning electron microscopy (SEM) images were acquired using a Zeiss SUPRA 40VP operating at an accelerating voltage of 3.0 kV. We analyzed the cross sections of screen-printed electrode, and nicotine sensor by SEM (SI Figure 23 and SI Figure 24). Distinct layers were shown. Additional thickness measurements are presented in SI Table 7.

### Materials

Phosphate buffered saline (PBS) was purchased from Life Technologies (Grand Island, NY). Iron(III) chloride, sodium chloride (NaCl), potassium chloride (KCl), calcium chloride (CaCl_2_), magnesium sulfate (MgSO_4_), sodium bicarbonate (NaHCO_3_), monosodium phosphate (NaH_2_PO_4_), sodium gluconate, glucose, sucrose, potassium ferricyanide(III), hydrochloric acid, chitosan, acetic acid, ethanol, bovine serum albumin (lyophilized powder, ≥96%), tetrahydrofuran, glutaraldehyde solution (Grade I, 8% in water), hydrogen peroxide solution, and Amicon® Ultra Centrifugal Filters, Whatman^®^ Grade 1 Filter Paper, benzoic acid, propylene glycol, and glycerol were purchased from Sigma-Aldrich (St. Louis, MO). Pierce Micro BCA Protein Assay kit, HisPur™ Ni-NTA Resin, Amplex® Red Hydrogen Peroxide/Peroxidase Assay Kit were purchased from Fisher Scientific (Pittsburgh, PA). Screen-Printed Electrodes TE100 were purchased from CH Instruments, Inc. (Austin, TX). Screen-printed carbon electrodes DRP-110, multi-walled carbon nanotubes modified screen-printed carbon electrode DRP-110CNT, screen-printed Prussian blue/carbon electrode DRP710 were purchased from Metrohm USA (Riverview, FL).

Artificial eccrine perspiration (artificial sweat) was purchased from Pickering Laboratories (Mountainview, CA). We obtained nonsmoker urine from a single volunteer donor, under an IRB approved procedure (IRB 803E). The collected urine was filtered with Whatman^®^ Grade 1 Filter Paper. An artificial interstitial fluid was prepared in water with 107.7 mM NaCl, 3.48 mM KCl, 1.53 mM CaCl_2_, 0.69 mM MgSO_4_, 26.2 mM NaHCO_3_, 1.67 mM NaH_2_PO_4_, 9.64 mM sodium gluconate, 5.55 mM glucose, and 7.6 mM sucrose^88^. Subsequently, PBS, artificial interstitial fluid, artificial sweat, and human urine were spiked with nicotine for generating dose response curves with optimized WT sensor.

### Strain Selection

*Pseudomonas putida* (Trevisan) Migula (ATCC^®^ BAA-2546^™^) was purchased from ATCC (Manassas, VA) and linked with a corresponding GenBank accession number (NC_015733.1). The strain is aerobic and was propagated and grown in ATCC^®^ Medium 18: Trypticase Soy Agar/Broth at 30°C as recommended by ATCC.

### Growth Curve

In order to determine the doubling time of the strain, growth curves were performed. All growth curves were done in 96 well flat clear bottom black polystyrene TC-treated microplates (Corning Inc., Corning, NY). Measurements were taken with an Infinite M200 Pro (TECAN Group Ltd., Medford, MA) spectrophotometer at 30°C. Readings were performed over 96 cycles of 15 minutes each at 600 nm absorbance with 25 flashes in a 3×3 (XY-Line) type reads per well. In between reads there was orbital shaking at 150 rpm frequency for a total of 10 minutes. To first characterize the growth alone, a ½ serial dilution of 8 concentrations from 0.5 – 0.0039 OD_600_ nm were prepared in M18 media. Then each concentration was measured as previously described by the TECAN microplate reader and normalized against a media background control in singlet. Afterward, an appropriate starting concentration of cells was chosen which shows a substantially long lag-phase, linear log phase, and a plateau at stationary-phase.

### Solvent Growth Curve

Once an appropriate starting cell concentration was chosen, a secondary growth curve was performed to test the toxicity levels of the solvent, H_2_O, used to dissolve nicotine. *P. putida S16* was incubated under microplate reader conditions previously mentioned at a starting OD determined by the first growth curve of 0.005 with H_2_O at ½ serial dilutions for a total of 8 concentrations (90 – 0.70%) tested in singlet. Two controls were included; a positive control without solvent and a media control which allowed for appropriate normalization. Solvent exposure growth curves allowed for choosing the maximum amount of solvent concentration that *P. putida S16* could sustain while maintaining relative viability in order to determine a range of nicotine concentrations which could be used.

### Nicotine Growth Curve

A tertiary growth curve was performed to test the toxicity levels of nicotine (Sigma-Aldrich Corp., St. Louis, MO) specific to *P. putida S16*. The strain was incubated under microplate reader conditions previously mentioned at a starting OD determined by the first growth curve and the highest solvent concentration with nicotine at ½ serial dilutions for a total of 8 concentrations tested in singlet. Three controls were included; a positive control with the chosen starting OD and the highest tolerable solvent concentration (%), a positive control with the chosen starting OD, and a media control alone which allowed for normalization. Nicotine exposure growth curves allowed for choosing the maximum amount of nicotine concentration that *P. putida S16* would sustain while maintaining relative phenotypic viability.

### Strain RNA Extraction

Cells were grown in 5 mL M18 at a starting OD_600_ of 0.005 with 21.6 µM nicotine diluted in H_2_O in 14 mL polypropylene round-bottom tubes (Corning Inc., Corning, NY) in singlet. The cells were incubated at 30°C with continuous orbital shaking at 150 rpm until the end of lag-phase (3 hours) and mid-log phase (7.5 hours) from the start of inoculation when RNA extraction was performed. Controls were grown in the same conditions without nicotine but in the same volume of H_2_O. At the time of RNA extraction, a 1:1 ratio of RNAprotect Bacteria Reagent (Qiagen Inc., Germantown, MD) was added followed by spinning down at 4°C for 10 minutes at 4000xg. Supernatant was removed and the pellet re-suspended in 300 µL of RNAprotect and transferred into 2.0 mL Safe-Lock Tubes (Eppendorf, Hauppauge, NY). The samples were then spun down at 4°C for 10 minutes at 10,000xg. RNA extraction was done by Qiacube (Qiagen Inc., Germantown, MD) set to the RNeasy Protect Bacteria Mini Kit protocol of bacterial cell pellet with enzymatic lysis. Lysis buffer was prepared as described by the protocol with the exception of the addition of 150 mg/mL lysozyme (Sigma-Aldrich Corp., St. Louis, MO) and 20 mg/mL proteinase K (F. Hoffmann-La Roche Ltd, Indianapolis, IN) all diluted in 1xTE buffer. RNA samples were subsequently quantified using Qubit RNA HS Assay Kit (Thermo Fisher Scientific Inc., Cambridge, MA) and analyzed using a RNA 6000 Pico Kit (Agilent Technologies Inc., Santa Clara, CA) in a 2100 Bioanalyzer (Agilent Technologies Inc., Santa Clara, CA). RNA samples were either immediately used for RNA-Seq library preparation or stored long-term at -80°C after addition of 1 µL RNase Inhibitor, Murine (New England Biolabs, MA).

### RNA-Seq Library Preparation

After RNA samples have been quantified and analyzed they were DNase treated using TURBO DNase 2 U/µL (Thermo Fisher Scientific Inc., Cambridge, MA) and cleaned using Agencourt RNAClean XP SPRI beads (Beckman Coulter, Inc., Brea, CA). RNA-Seq libraries were then produced from these samples using a modified ScriptSeq v2 RNA-Seq Library Preparation Kit (Illumina Inc., San Diego, CA) ensuring use of unique index primers through ScriptSeq Index PCR Primers (Sets 1-4) 48 rxns/set (Illumina Inc., San Diego, CA). Libraries were quantified by both a Qubit dsDNA HS Assay Kit (Thermo Fisher Scientific Inc., Cambridge, MA) and by Bioanalyzer with the High Sensitivity DNA Kit (Agilent Technologies Inc., Santa Clara, CA). The samples were then pooled to 2 nM and submitted to the Boston University Microarray and Sequencing Resource Core Facility. Whole transcriptome RNA sequencing was performed by a NextSeq 500 (Illumina Inc., San Diego, CA) at high output (400 M reads) with 75 bp paired end sequencing read length. The data was then analyzed in-house through a Galagan Lab computational pipeline using DESeq for differential expression analysis.

### RNA-Seq Data Analysis

Adapter sequences were removed from reads and low quality bases trimmed from both ends using Cutadapt^89^. Reads were aligned to the reference genome with Bowtie2^90^. BAM files were sorted and indexed using Samtools^91^. Transcript assembly and expression quantification was performed using Cufflinks^92^. All resulting raw expression counts were normalized as a group using deseq^93^. A custom MATLAB script was then used to calculate fold changes of normalized counts for each gene between each nicotine exposure experiment and its corresponding solvent control. Genes +/- 1 log2 fold change from between the difference of nicotine vs a solvent control was deemed differentially expressed.

### Enzyme Expression and Purification

WT NicA2 and variants were recombinantly produced with poly-histidine tags in *E. coli* per previous work^1,55,58^. Plasmids were chemically transformed into *E. coli* BL21 (DE3) (New England Biolabs, MA) and purified as previously described^1^ with induction of lysogeny broth cultures at OD_600_ 0.6 to 0.8 using 1 mL of 1M isopropyl β-D-1-thiogalactopyranoside (IPTG) before analysis by SDS-polysaccharide gel electrophoresis (SDS-PAGE). Confirmed expression prompted 1 L cultures for expression and protein purification through disruption of cells using lysozyme and by passing cell lysate over a HisPur™ Ni-NTA Resin-packed column (Fisher Scientific, Pittsburgh, PA). Final protein products were quantified using the Pierce Micro BCA Protein Assay kit (Fisher Scientific, Pittsburgh, PA).

### *In Vitro* Enzyme Characterization

Amplex UltraRed assays were performed per instruction of the manufacturer (Thermo Fisher Scientific, Cambridge, MA). Final concentrations of Amplex UltraRed, 500 µM, and HRP, 1 U/mL, in a total volume of 20 µL were used. NicA2 was washed and concentrated 3x with H_2_O using an Amicon® Ultra 0.5 mL Centrifugal Filter (MilliporeSigma, Burlington, MA). The enzyme concentration was measured using a Nanodrop 2000 Spectrophotometer (Thermo Fisher Scientific Inc., Cambridge, MA). All measurements of fluorescence were done in a 384 well flat bottom black polystyrene microplate (Corning Inc., Corning, NY) and read in an Infinite M200 Pro (TECAN Group Ltd., Medford, MA) spectrophotometer at room temperature with the excitation set to 490 nm and emission at 585 nm. Readings were done over 1 hour with each well read every 30 seconds.

### Evaluation of immobilization methods

A Prussian Blue (PB) mediator layer was deposited onto TE100 SPEs by cyclic voltammetry. PB solution was prepared by mixing equal volume of 10 mM FeCl_3_, 400 mM KCl, 10 mM K_3_Fe(CN)_6_, and 400 mM HCl. The electrodes were dipped into the PB solution and the potential was swept from 0 to 0.5 V (versus Ag/AgCl) for 1 cycle at a scan rate of 0.02 V/s.

To select the optimal immobilization method, we evaluated four different immobilization strategies for NicA2 on Prussian Blue deposited screen-printed electrodes (PB/TE100). Immobilization method 1: equal volume of 400 µM NicA2 and 1 wt% chitosan in acetic acid (2.0,1.5,1.0, 0.5 wt%, respectively) were mixed. 10 µL above solution was casted on the PB/TE100, kept in fridge (4 °C) till dry. Immobilization method 2: 5.0 µL of 400 µM NicA2 solution in PBS was casted onto the PB/TE100 surface. The electrodes were dried at room temperature (25 °C). Subsequently, 20 µL of 1% Nafion (in 95% ethanol) was dropped onto the electrode surface and dried. The electrodes were stored in the refrigerator when not in use. Immobilization method 3: 400 µM NicA2 solution (containing 10 mg/mL bovine serum albumin as stabilizer) was mixed with chitosan solution (1 wt% chitosan in 0.5 wt% acetic acid) in a 1:1 v/v ratio. Subsequently, a 10 µL droplet of the above solution was casted on the PB/TE100 and dried under ambient conditions. Thereafter, 5 µL of polyvinyl chloride (PVC) solution (3 wt% in tetrahydrofuran) was drop casted and allowed to dry under ambient conditions for at least 3 h before use. Immobilization method 4: 400 µM NicA2 solution (containing 10 mg/mL bovine serum albumin as stabilizer) was mixed with chitosan solution (1 wt% chitosan in 0.5 wt% acetic acid) in a 1:1 v/v ratio. Subsequently, a 10 µL droplet of the above solution was casted on the PB/TE100 and dried under ambient conditions. Thereafter, 10 µL 2 wt% glutaraldehyde solution was drop-casted on the sensor for robust crosslinking of the enzyme layer.

For nicotine detection, 20 µL PBS was added to cover the NicA2 coated electrodes. With a voltage of -0.2 V applied using the VersaSTAT MC potentiostat, the time and current were recorded. At a predefined time point, 20 µL 400 µM nicotine in PBS was added and the current was measured as a function of time. The four different immobilization strategies were compared.

### Screen printed electrode comparison

To select the optimal screen-printed electrode, we evaluated four different types of SPEs for hydrogen peroxide detection: Screen-Printed Electrodes TE100 were purchased from CH Instruments, Inc. (Austin, TX). Screen-printed carbon electrodes DRP-110, multi-walled carbon nanotubes modified screen-printed carbon electrode DRP-110CNT, screen-printed Prussian blue/carbon electrode DRP710 were purchased from Metrohm USA (Riverview, FL). TE100 possesses a carbon working electrode (diameter=3 mm), a carbon counter electrode, and an Ag/AgCl pellet reference electrode. DRP-110 possesses a carbon working electrode (diameter=4 mm), a carbon counter electrode, and a silver reference electrode.DRP-110CNT possesses a multi-walled carbon nanotubes/carbon working electrode (diameter=4 mm), a carbon counter electrode, and a silver reference electrode. DRP-710 possesses a Prussian blue/carbon working electrode (diameter=4 mm), a carbon counter electrode, and a silver reference electrode. Compared to TE100’s graphite lead terminal and smaller active surface, DRP-110, DRP-110CNT, and DRP-710 possess silver lead terminal and larger rough active surface. Both features have been previously associated with elevated sensitivity due to better metal connectivity with measurement set up and increased effective electrode active surface area.TE100, DRP-110, and DRP-110CNT were electrodeposited with different thickness of PB by cyclic voltammetry. For thin layer of PB, the electrodes were dipped into the PB solution and the potential was swept from 0 to 0.5 V (versus Ag/AgCl) for 1 cycle at a scan rate of 0.02 V/s. For medium layer of PB, the electrodes were dipped into the PB solution and the potential was swept from -0.5 V to 0.6 V (versus Ag/AgCl) for 5 cycle at a scan rate of 0.05 V/s. For thick layer of PB, the electrodes were dipped into the PB solution and the potential was swept from -0.5 V to 0.6 V (versus Ag/AgCl) for 10 cycle at a scan rate of 0.05 V/s. DRP-710 electrodes, pre-impregnated with Prussian Blue, were used as received without further PB deposition. Hydrogen peroxide detection capability was evaluated by monitoring the current generated upon addition of hydrogen peroxide solution in PBS, with a voltage of -0.2 V applied. Selection was predicated upon electrode eliciting the highest current.

### Dose response curves in PBS with varying enzyme quantity

After selection of optimal immobilization method and screen-printed electrode, we established dose-response curves for nicotine detection in PBS with varying enzyme quantity. Equal volume of NicA2 solution and 1 wt% chitosan in 0.5 wt% acetic acid were mixed. 10 µL above solution was casted on the DRP-710, kept in fridge (4°C) till dry. Four concentrations of NicA2 solution were used: 200 µM, 400 µM, 800 µM, and 1600 µM, rendering 1,2,4, and 8 nmol NicA2 on the electrodes, respectively. The enzyme was concentrated with Amicon® Ultra Centrifugal Filters to desired concentration. Chronoamperometric response of the nicotine sensor with different enzyme amounts were carried out at potential -0.2 V (versus Ag/AgCl) with VersaSTAT MC. The sensor was calibrated in the nicotine concentration range between 0 and 1 mM.

### Dose response curves in complex matrices with optimized WT sensor

We selected the following configuration for our final WT NicA2 biosensor: immobilization with 1 wt% chitosan in 0.5 wt% acetic acid; DRP-710 electrode; 2 nmol deposited NicA2. We established dose-response curves for nicotine detection in additional fluids: interstitial fluid, sweat, and urine. Chronoamperometric response of the nicotine sensor were carried out at potential -0.2 V (versus Ag/AgCl) with VersaSTAT MC. The sensor was tested in the nicotine concentration range between 0 and 200 µM.

### Biosensor sensitivity calculations

The LOD is defined as the nicotine concentration yielding a signal greater than 3 times the pool standard deviations above background. LOQ is defined as the nicotine concentration yielding a signal greater than 10 times the pool standard deviations above background

### Quantification of nicotine in e-cigarette liquid

To further validate the applicability of the sensor, we subjected the sensor to detect nicotine in e-cigarette liquid. We obtained Juul^®^ pod, which contains 5% (w/w) nicotine, from a local smoke shop. To evaluate the selectivity of the sensor, chronoamperometric responses to nicotine were measured in the presence of the common interferants (benzoic acid, propylene glycol, and glycerol) in e-cigarette liquid. Addition of 30 µL 200 µM nicotine solution to 30 µL mixture of benzoic acid, propylene glycol, and glycerol was performed at predefined time point and the current was recorded over time. Similar experiments were performed to evaluate the sensor response to interferants. Current was recorded where 30 µL interferants solution was added to 30 µL PBS. After evaluating the selectivity of the sensor, the calibration curve was established with 0, 25, 75, 150, 250 μM nicotine in PBS with the addition of benzoic acid, propylene glycol, and glycerol to mimic the e-cigarette liquid composition. 10 mM nicotine was prepared by adding 81.1 mg nicotine, 61.6 mg benzoic acid, 443.9 mg propylene glycol, 1035.7 mg glycerol to 50 mL PBS^94^. 25, 75, 150, and 250 μM nicotine were prepared by diluting above 10 mM nicotine solutions with PBS. Additionally, the e-cigarette liquid from Juul^®^ pod was diluted to 50, 100, and 200 μM with PBS. The quantification of nicotine was carried out by chronoamperometry, and the recovery of nicotine was calculated.

### Biosensor Preparation

Electrochemical experiments were performed with a DropSens µSTAT 4000P (Metrohm USA, Riverview, FL). Sensors were prepared with a DS710 Prussian Blue/Carbon Electrode (Metrohm USA, Riverview, FL) or a custom made DS710BIG Prussian Blue/Carbon Electrode (Metrohm USA, Riverview, FL). Prior to enzyme deposition, Equal volume of NicA2 (400 µM) and 1 wt% chitosan in 0.5 wt % acetic acid (Sigma-Aldrich Corp., St. Louis, MO) were mixed. NicA2 was immobilized onto the SPEs by drop-casting 10 µL of this mixture onto the working electrode. 40 µL of this mixture were used for preparing the DS710BIG. The sensors were allowed to dry overnight at room temperature prior to use.

### Biosensor Characterization by Pipetting

All chronoamperometric experiments were performed at a potential of -0.2 V (versus RE). At the beginning of the experiment, 40 µL H_2_O were added to connect all three electrodes. Analyte additions were added at a defined volume, concentration, and time depending on the experiment being performed. Current generated was measured by integrating between a straight line from the end of the current response to the start and the signal itself.

### Biosensor Human Bodily Fluids Accuracy Study

Measurement of blindly doped human bodily fluids were done by first calibrating the biosensor with three known nicotine concentrations diluted in H_2_O. Then the bodily fluid, non-doped, was added, followed by three blindly doped samples. Artificial eccrine perspiration (Pickering Laboratories #1700-0022, Mountainview, CA) and artificial urine (Pickering Laboratories #1700-0558, Mountainview, CA) were used for testing. In compliance with Boston University institutional review board (IRB) approval (protocol #4803E) human non-smoker urine and sweat were collected for doping with nicotine and testing. Once the data was collected a calibration curve was generated from the three known nicotine concentrations and used to measure the blindly doped non-smoker samples.

### WearStat Construction

The WearStat was designed as a standalone wearable potentiostat with Bluetooth low energy (BLE) communication, paper-based fluidics, and a connector specific to the SPEs described herein. The four main subsystems of the device include the electronics, the SPE, the fluidics, and the 3D-printed enclosure. The electronics are composed of a custom analog printed circuit board (PCB) that is soldered on top of a small commercial clone of the Arduino Uno, DFRobot’s Beetle BLE, which serves as the microcontroller of the device and carries out BLE communication with a PC. The WearStat fluidic component is housed by two layers of PDMS that make up its main structure and hold the paper channel and sink over the SPE. This material was chosen due to its flexibility and its compatibility with rapid prototyping using 3D printed molds. The paper fluidic channel was designed by using Solidworks and cut to specific dimensions using a Zing 16 laser cutter (Epilog Laser, Mississauga, ON). The fluidic channel is composed of a combination of two different blotting papers GE Whatman GB003 (MilliporeSigma, Burlington, MA) and Ahlstrom 320 (Fisher Scientific, Pittsburgh, PA). The thinner paper remains in contact with both the skin and the surface of the SPE so that sweat is carried to the sensing components quickly as it is generated by the subject. The thicker, higher capacity paper is placed at the end of the system and acts as a sink to draw in sweat after measurements have been taken and maintain continuous flow through the device. A MATLAB GUI was created to interface with the WearStat to set the applied potential, choose a current range, record the current, and export the data.

### WearStat Flow Experiments

Multiple Miracle Sponge (The Color Wheel Company, Philomath, OR) sponges cut into 1 in^2^ were saturated with 20 mL of known concentrations of analyte were prepared and placed into separate petri dishes, which simulated a sweat-saturated skin surface. The device was put in contact with the sponges whereupon the paper channel wicked the solution from the sponge to the SPE where current was measured and transmitted to a PC using BLE in real-time. The device was then moved between sponges and a step response signal observed which corresponds to the analyte concentration. A syringe pump NE-300 (New Era Pump Systems, Inc. Farmingdale, NY) was used to control the flow rate (µL/m) of analytes of known concentrations in disposable syringes.

### Statistical analysis

Unless otherwise indicated, data presented were generated from three independent experiments. Error bars represent standard deviation values. Multivariance analysis was done using one-way ANOVA that was corrected using the Bartlett’s variance test, and for multiple comparisons, the Bonferroni multiple-comparison test was used. Statistical analysis was performed using the GraphPad Prism software (version 7.0; GraphPad Software Inc.) or MATLAB software.

## Supplemental Information [Figures]

**SI Figure 1.**
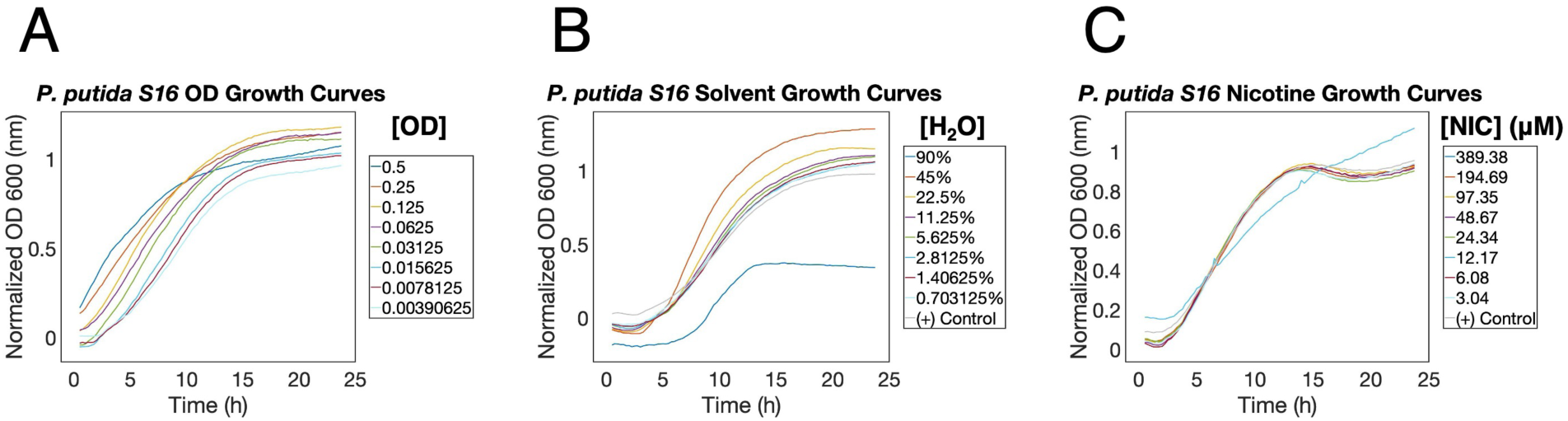
*P. putida S16* growth curves were performed to choose the optimal experimental conditions for RNA-Seq. **(A)** An OD growth curve with a ½ serial dilution found that starting ODs of 0.015625 and under provide a visible lag-phase. **(B)** A solvent growth curve with a ½ serial dilution found that the solvent used to dilute nicotine, H_2_O, does not affect *P. putida S16* physiology significantly under 11.25% final concentration. **(C)** A nicotine growth curve with a ½ serial dilution found that the full range of final [nicotine] tested do not affect *P. putida S16* physiology as compared to control.

**SI Figure 2.**
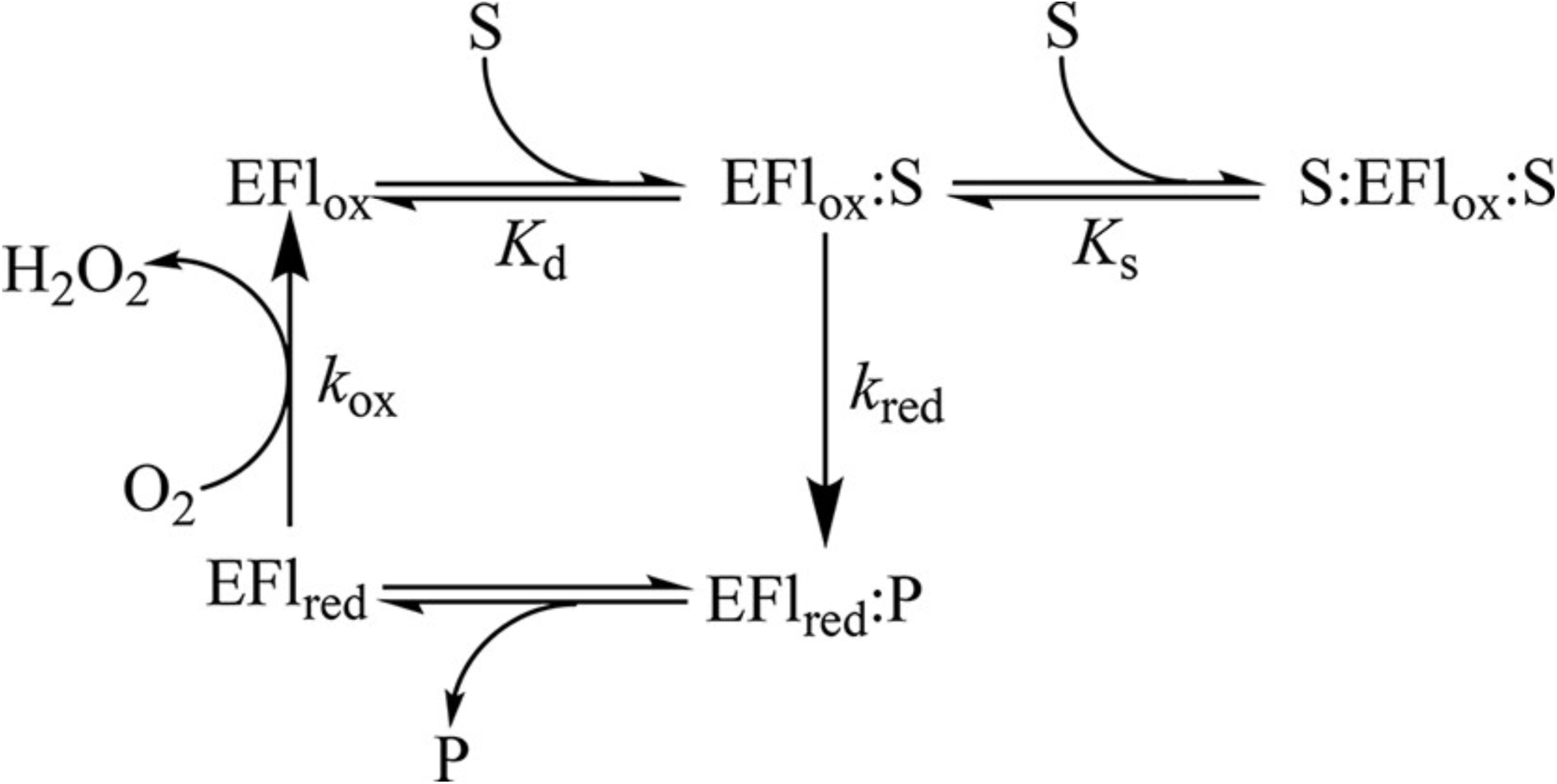
NicA2 mechanism^1^. Enzyme flavin (EFl), substrate (S), product (P).

**SI Figure 3.**
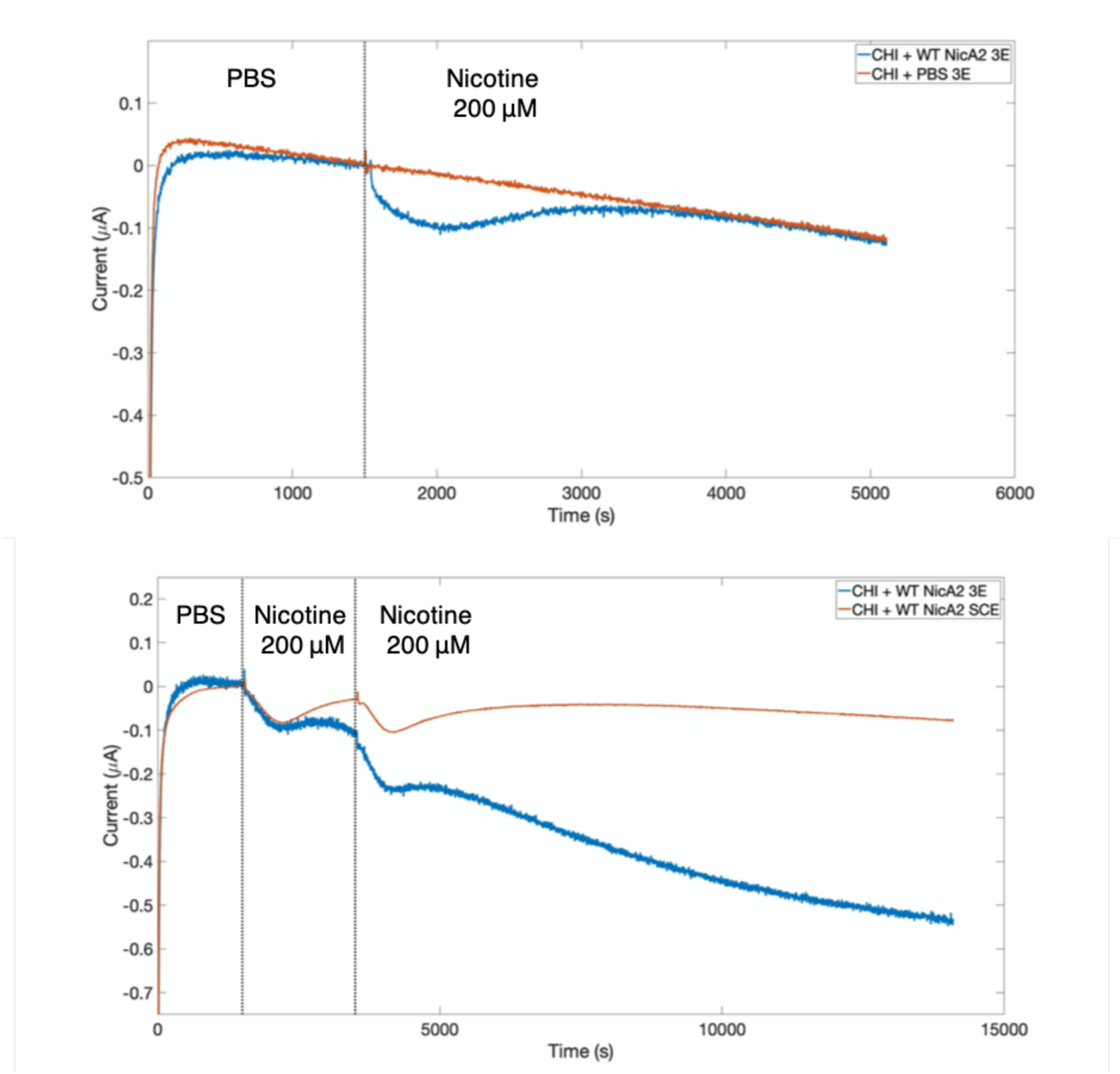
A two electrode (shorted counter electrode, SCE) vs a three electrode (3E) configuration results in less drift and noise.

**SI Figure 4.**
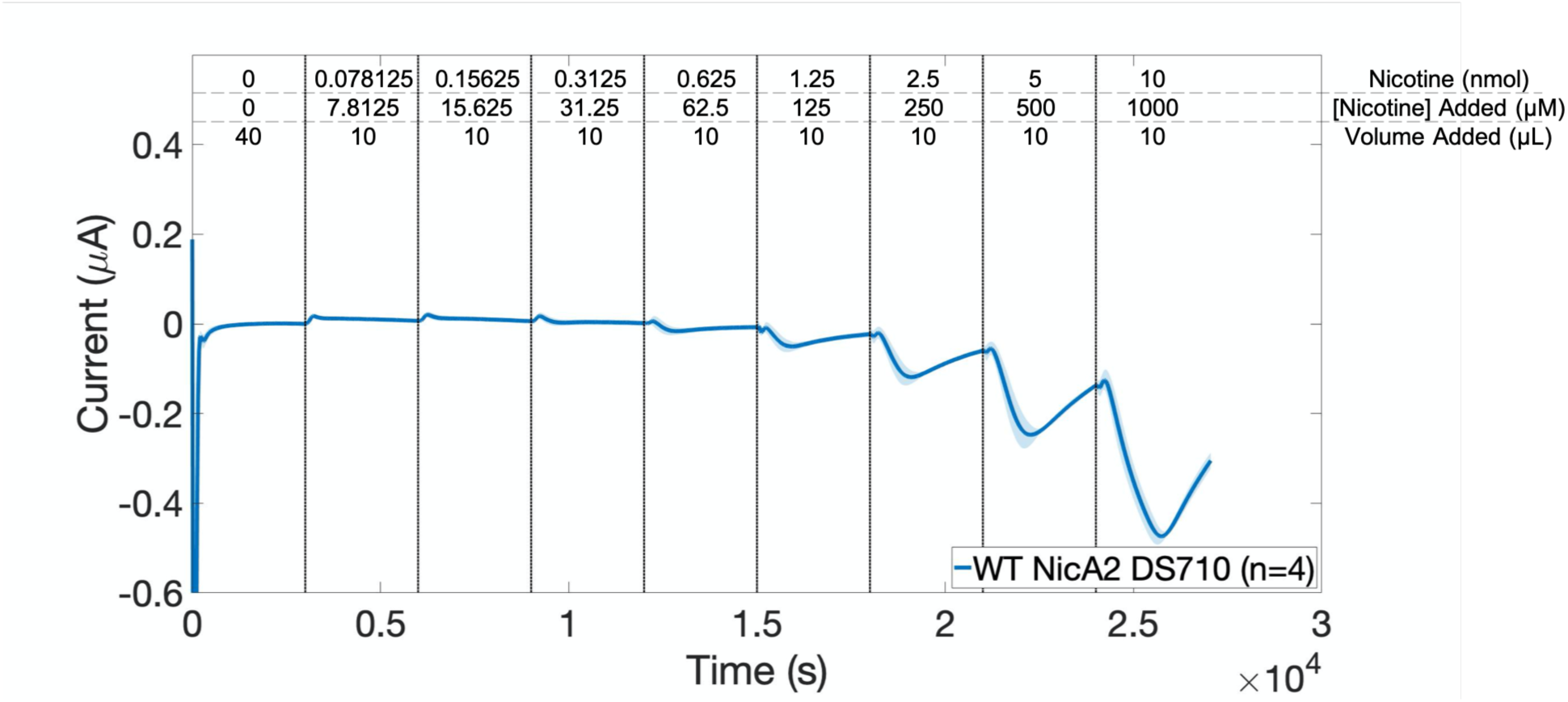
WT NicA2 DS710 dose response curve.

**SI Figure 5.**
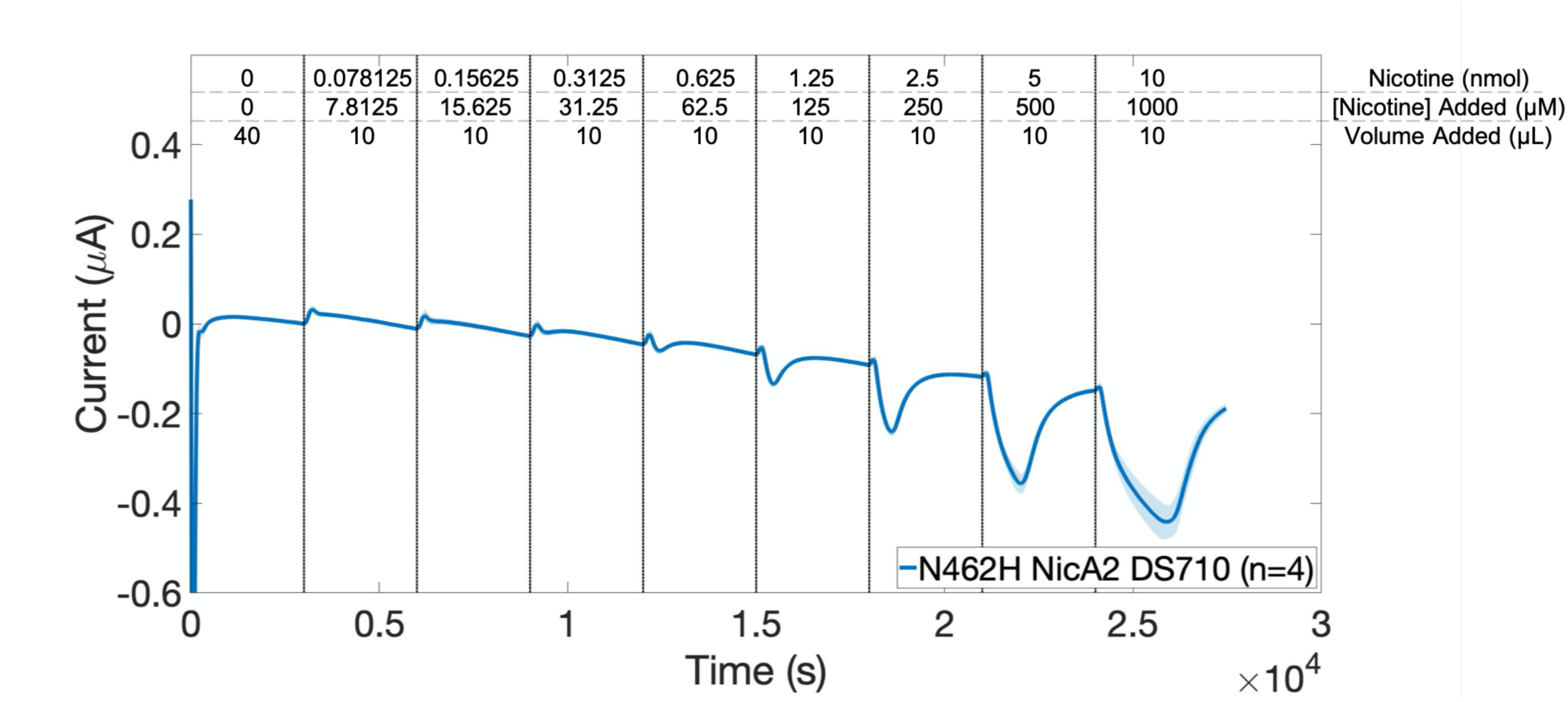
N462H NicA2 DS710 dose response curve.

**SI Figure 6.**
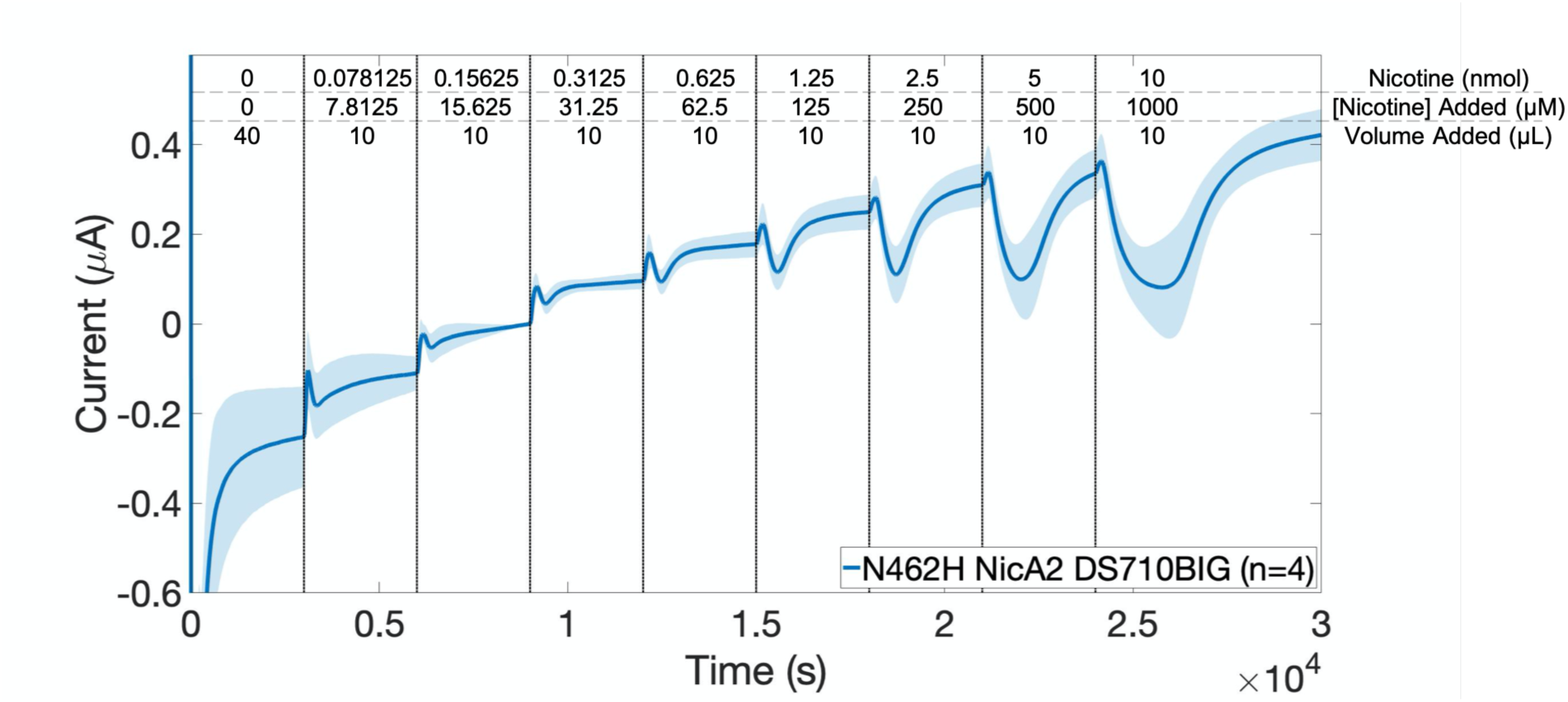
N462H NicA2 DS710BIG dose response curve.

**SI Figure 7.**
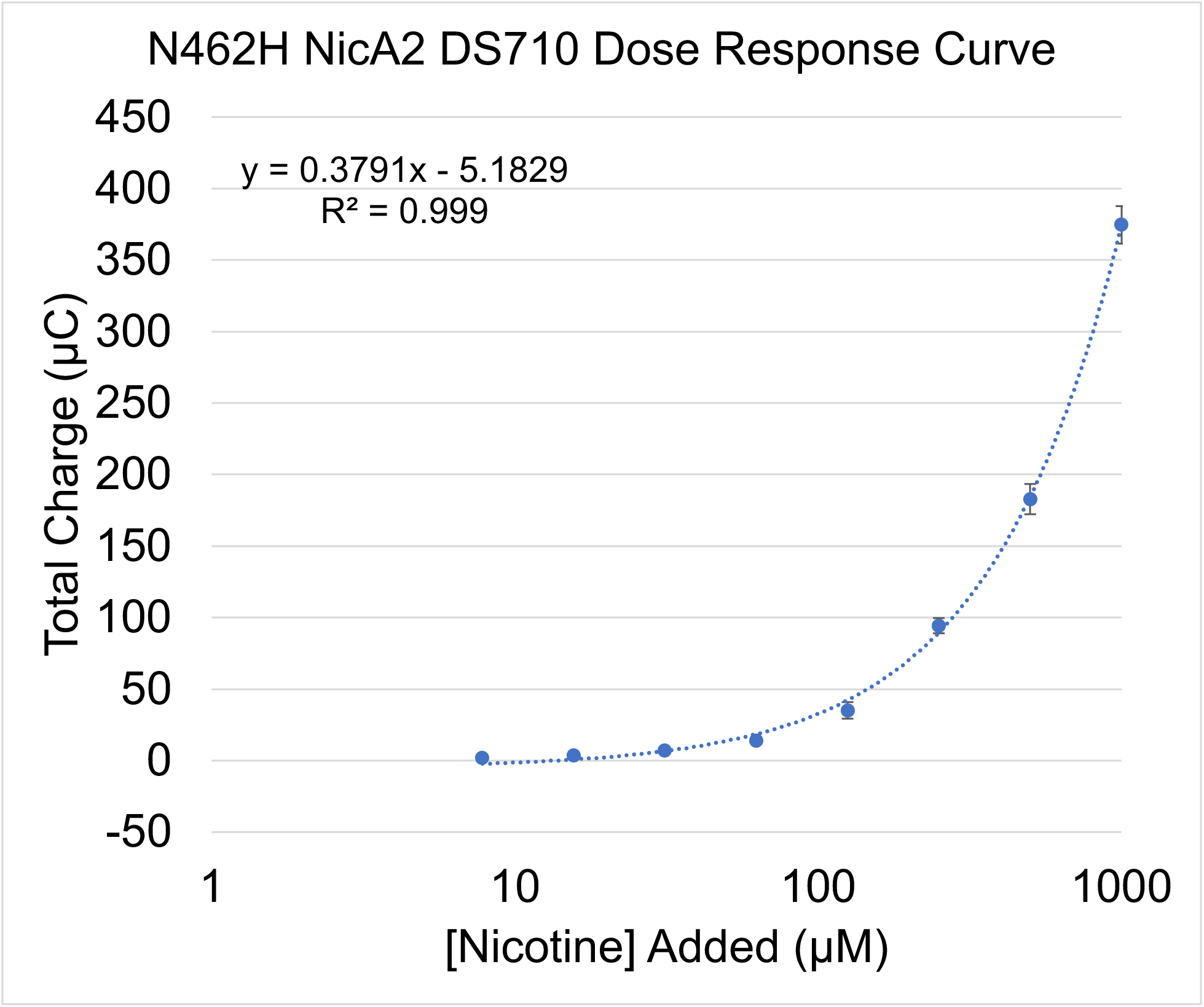
N462H NicA2 DS710 dose response curve and linear regression.

**SI Figure 8.**
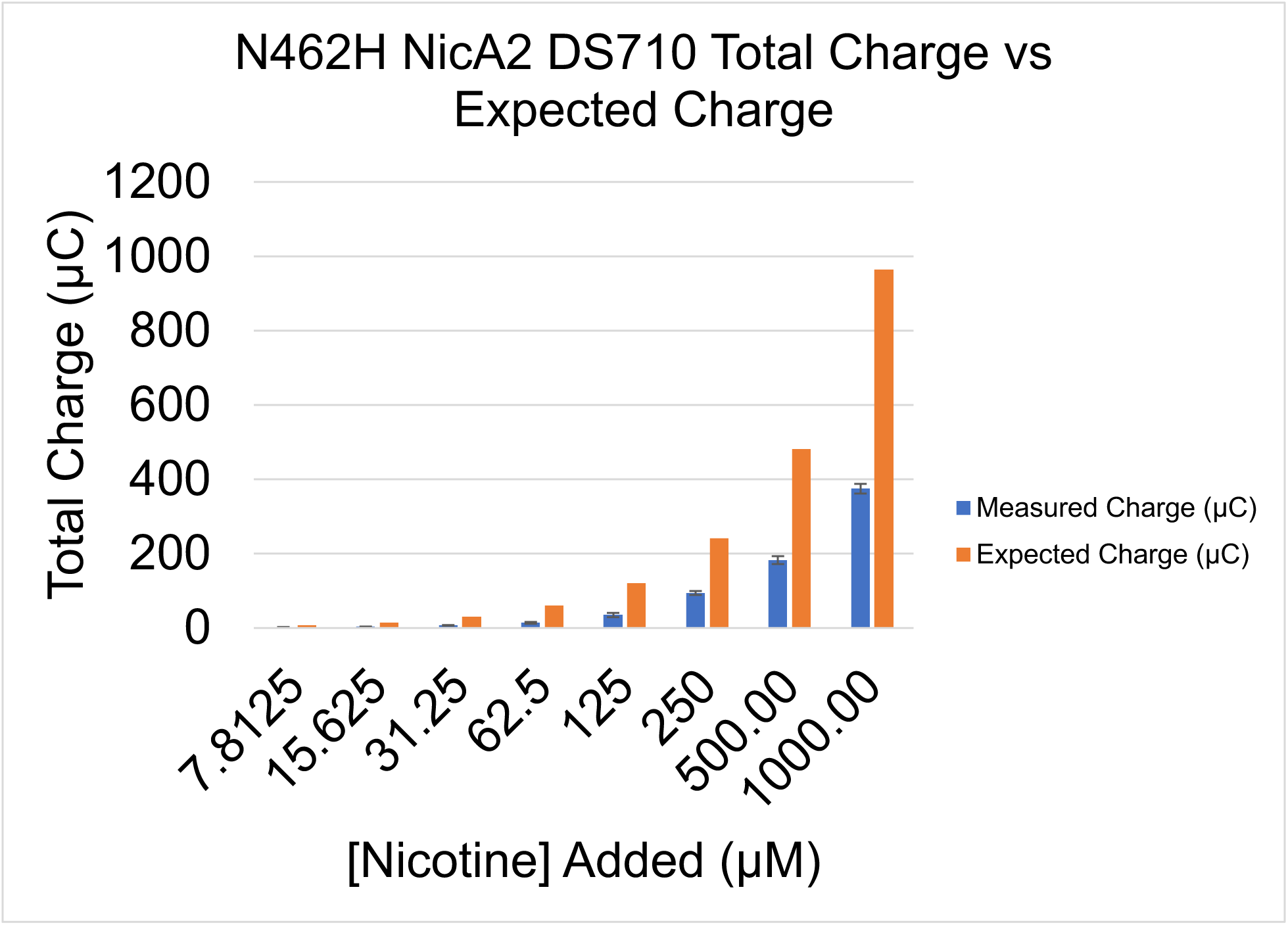
N462H NicA2 DS710 dose response total charge vs expected charge shows that not all of the nicotine in a droplet diffuses to the electrode surface.

**SI Figure 9.**
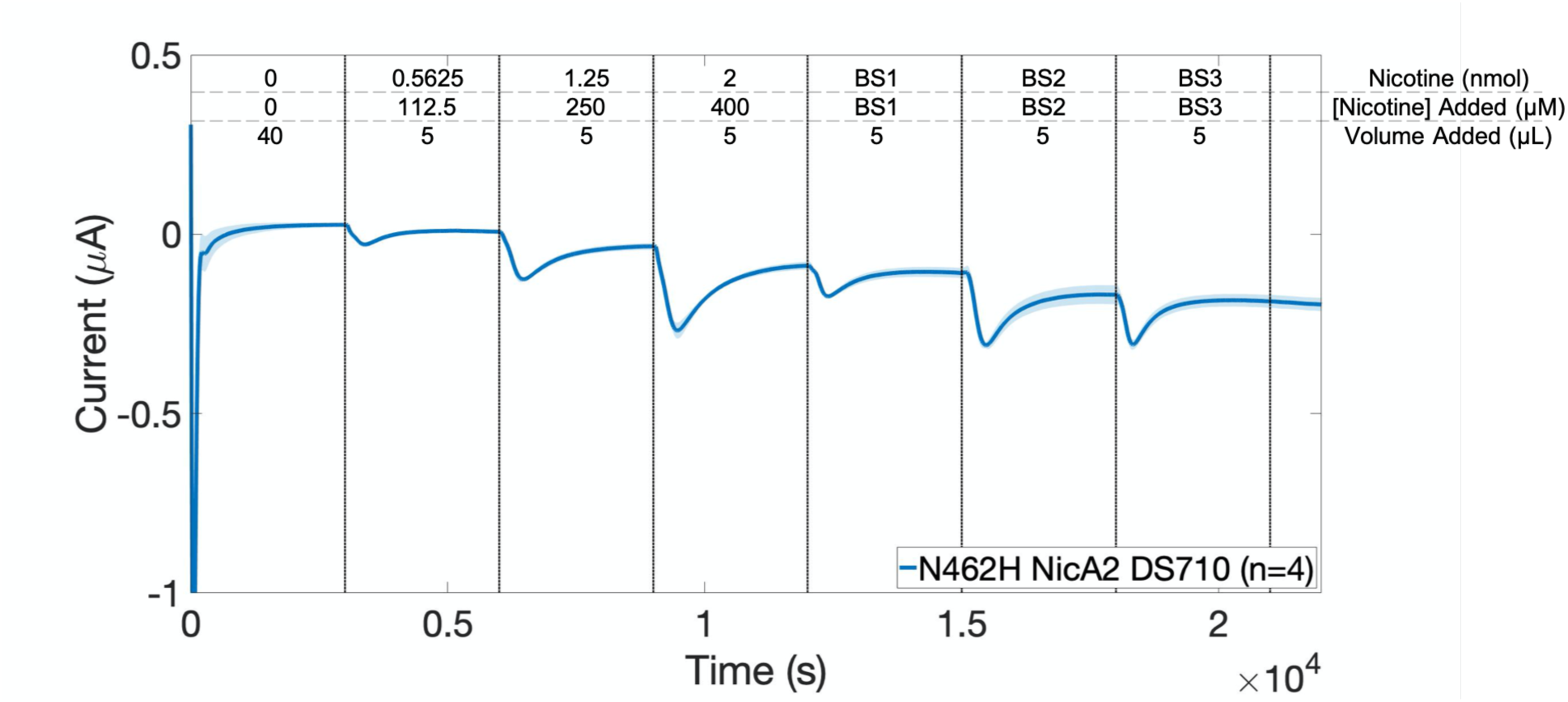
N462H NicA2 DS710 blind, doped PBS sample measurement.

**SI Figure 11.**
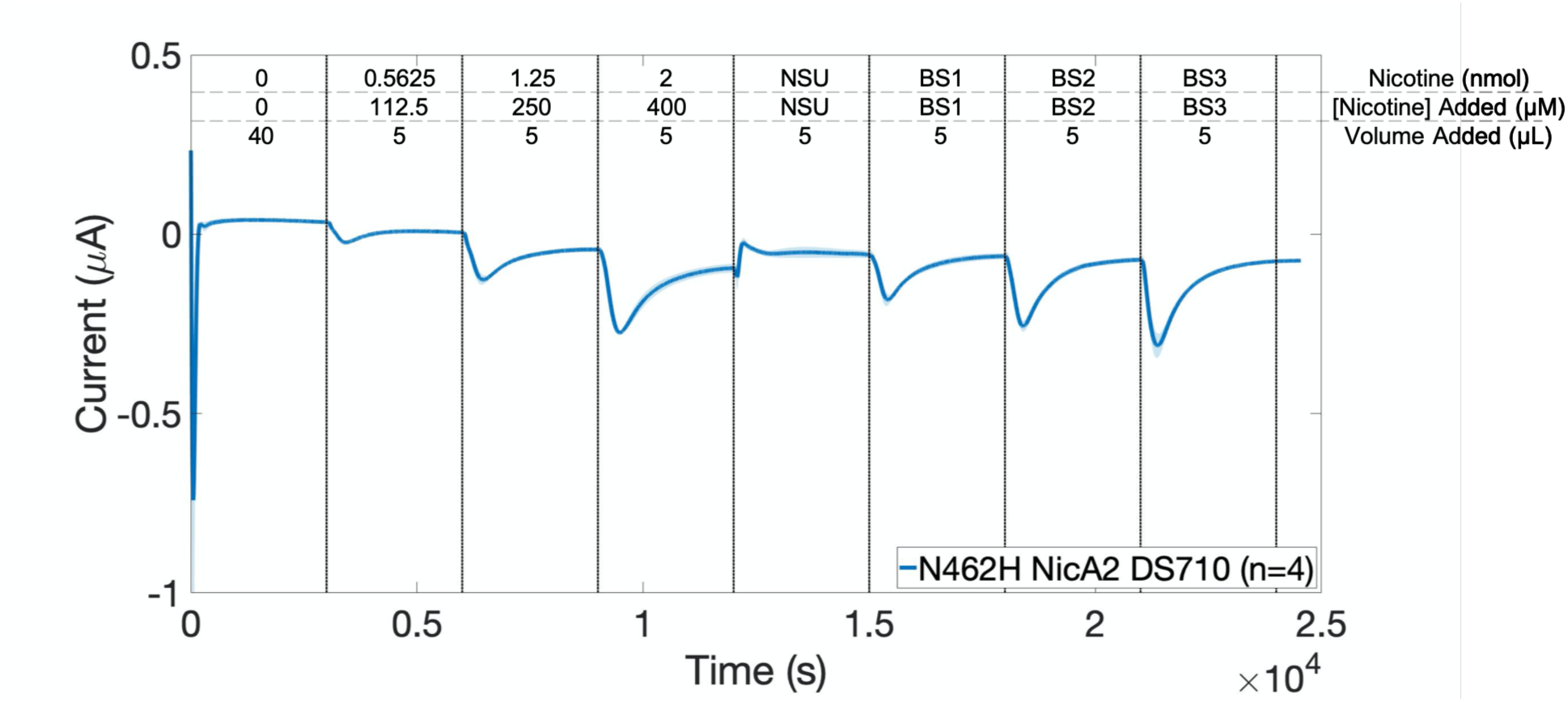
N462H NicA2 DS710 blind, doped non-smoker urine sample measurement.

**SI Figure 12.**
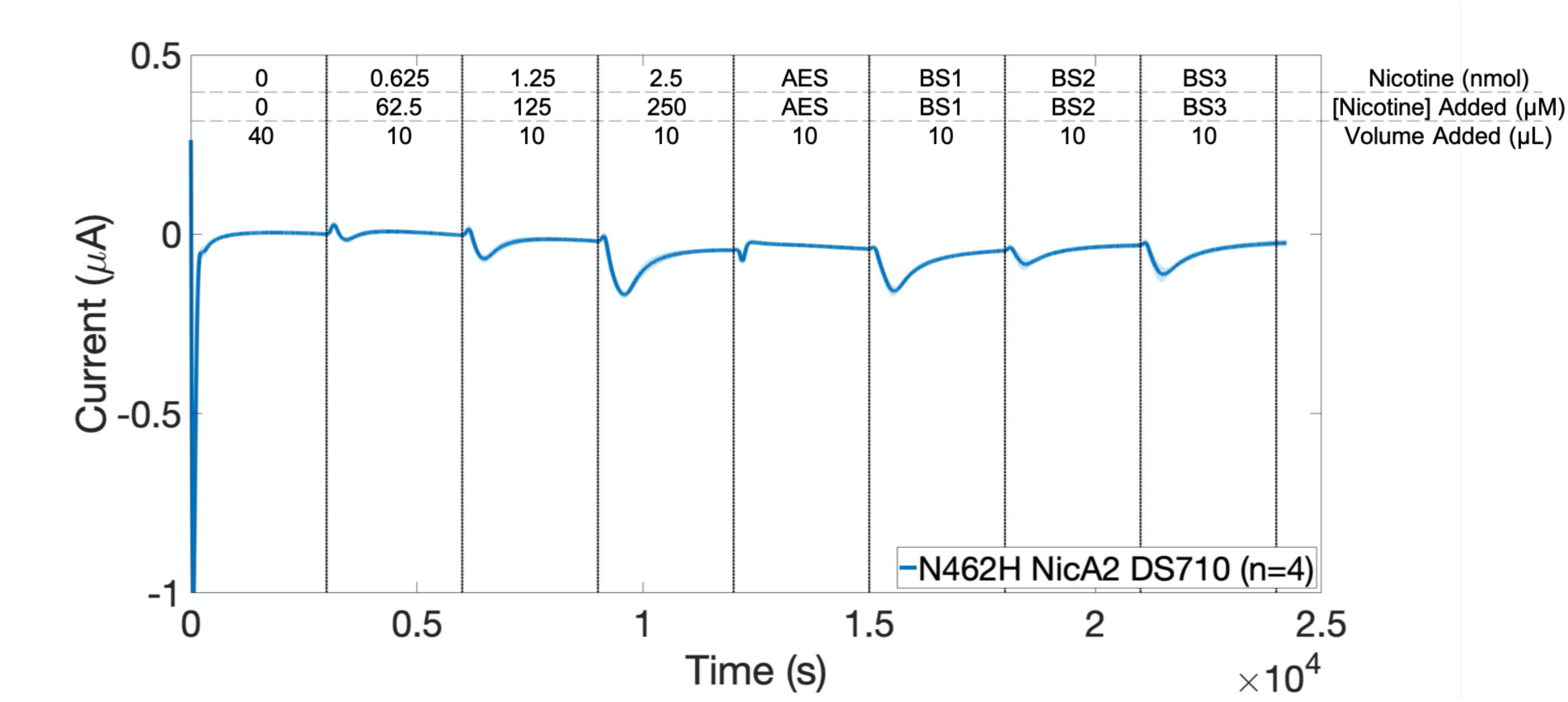
N462H NicA2 DS710 blind, doped artificial sweat sample measurement.

**SI Figure 13.**
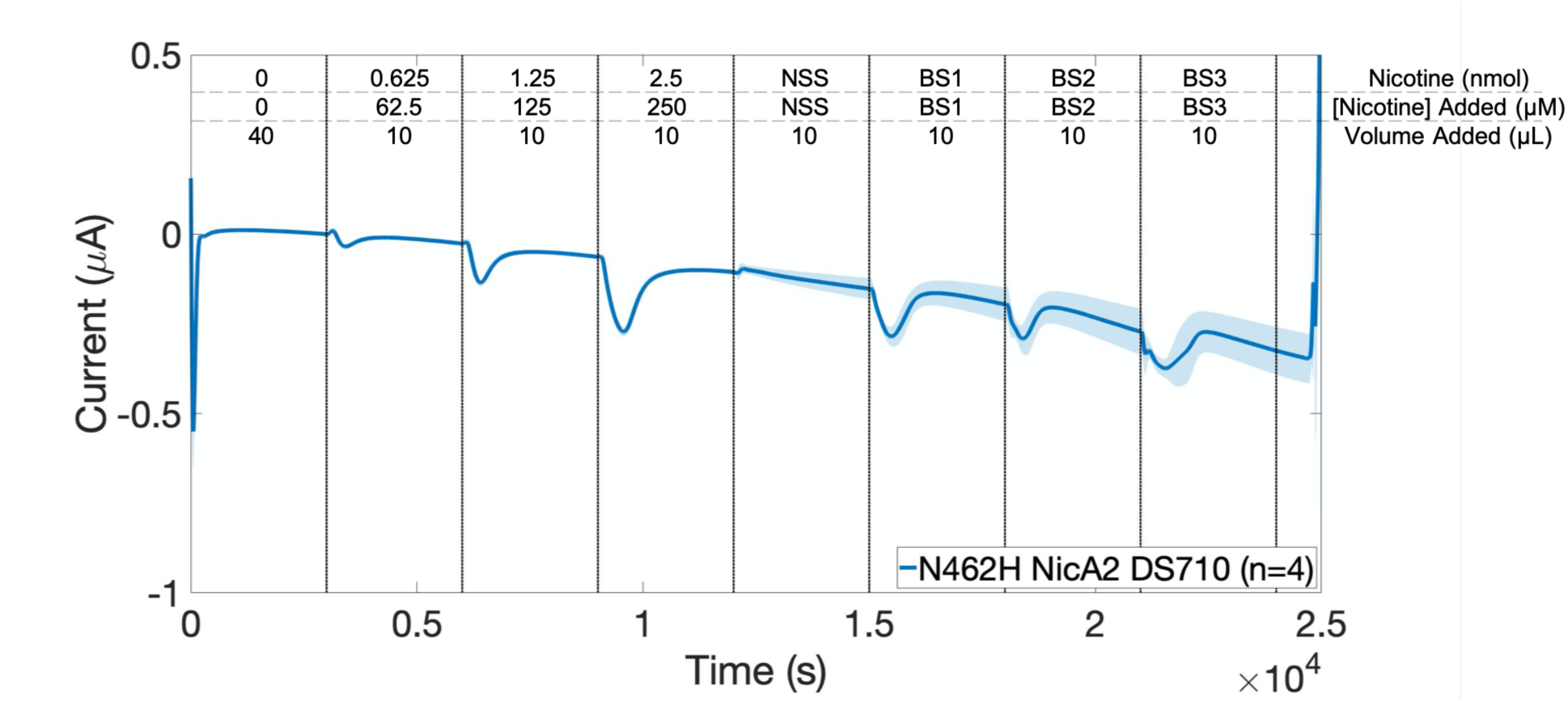
N462H NicA2 DS710 blind, doped non-smoker sweat sample measurement.

**SI Figure 14.**
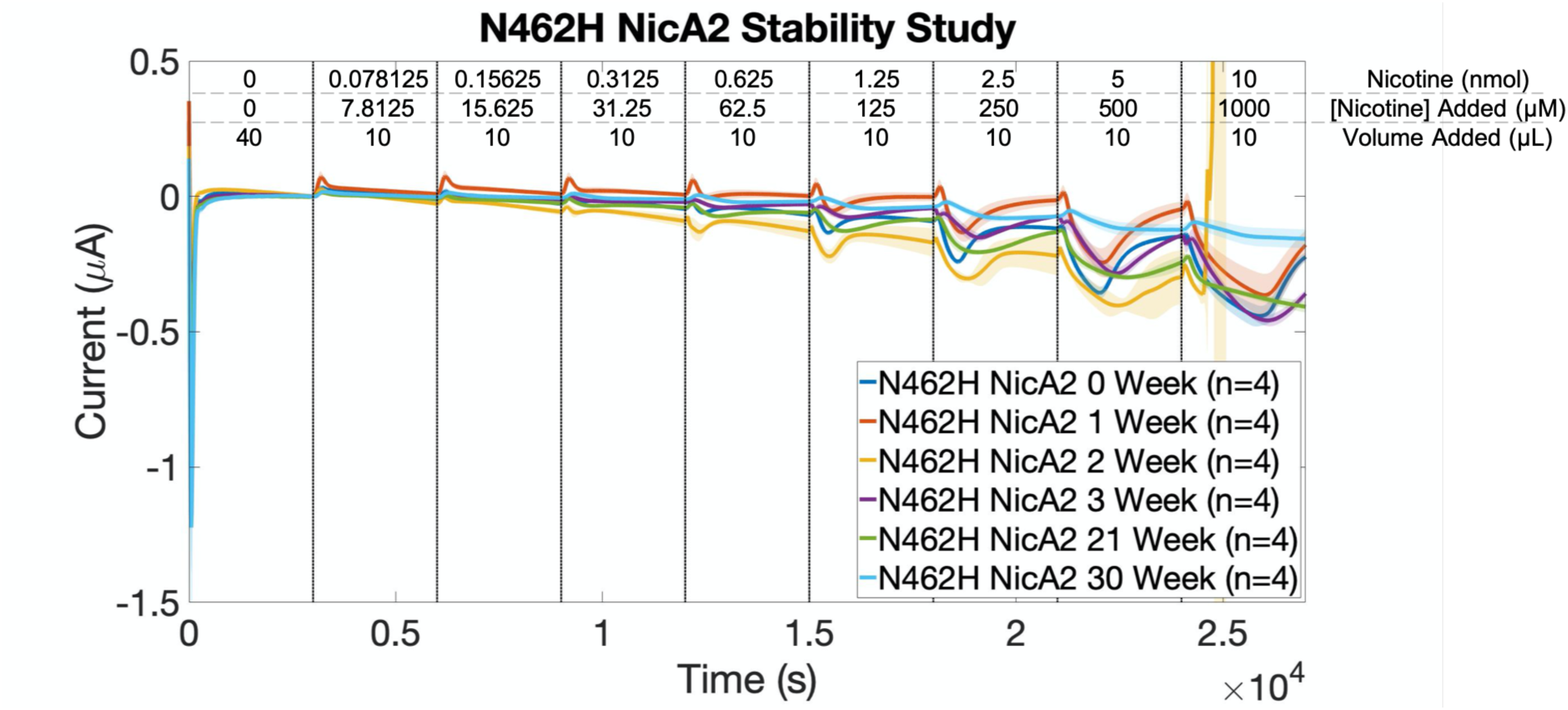
N462H NicA2 DS710 stability study with SPE storage at room temperature over the noted number of weeks.

**SI Figure 15.**
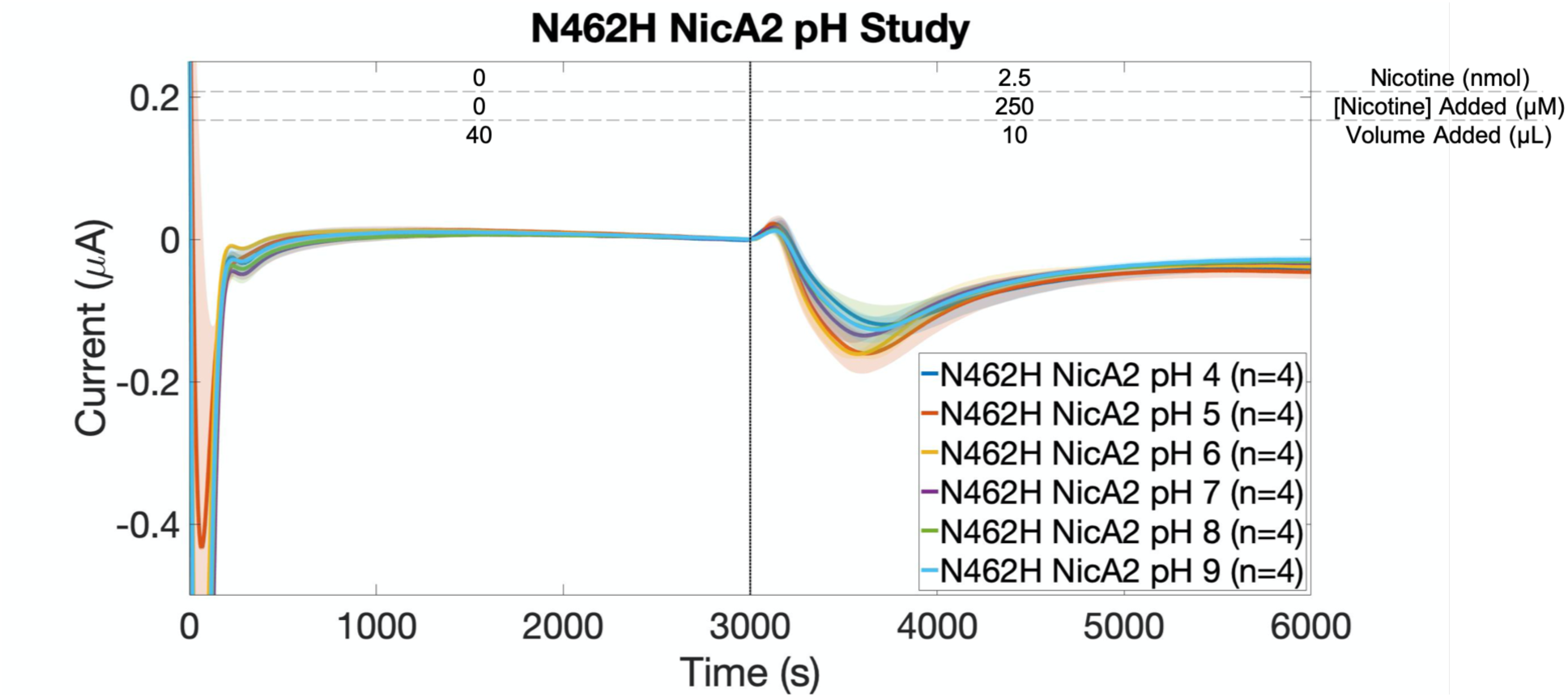
N462H NicA2 DS710 pH study showed similar activity across the entire pH range of 4-9.

**SI Figure 16.**
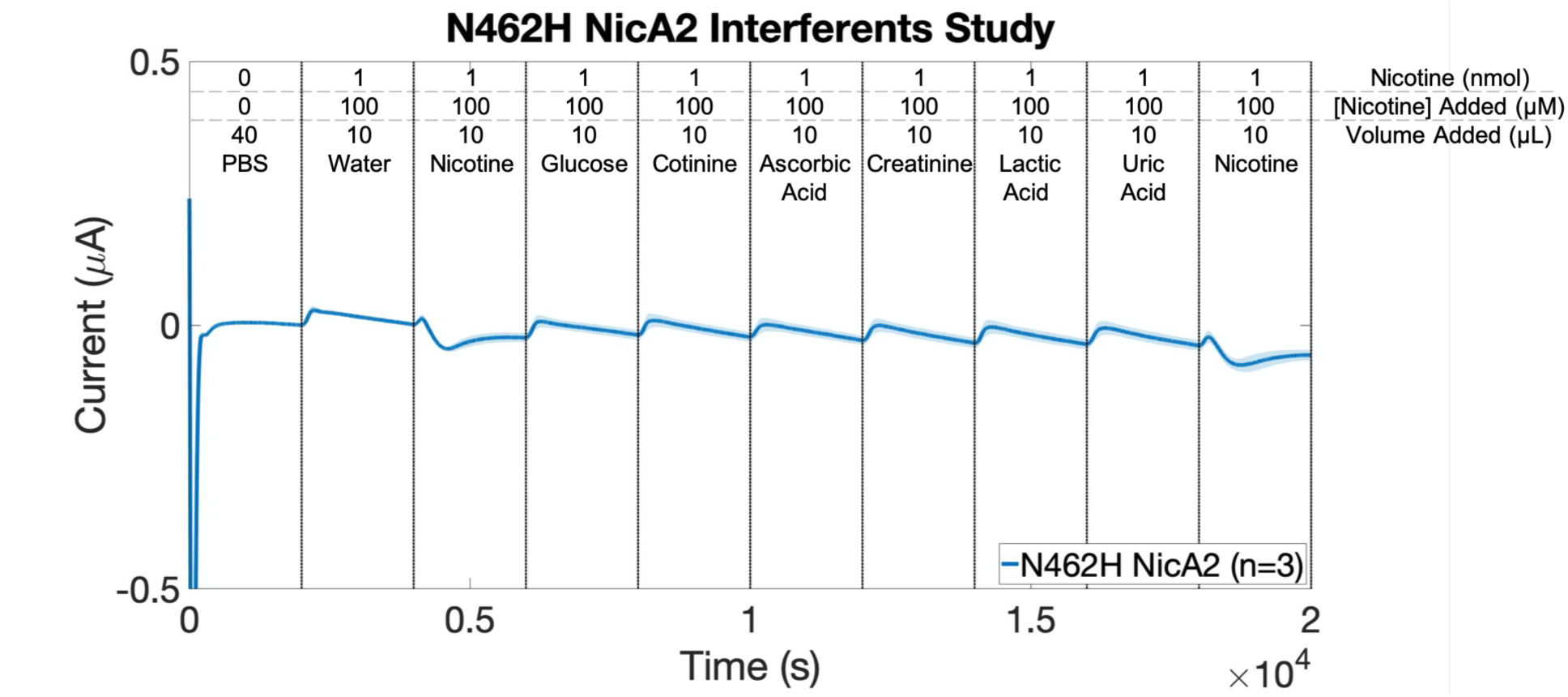
N462H NicA2 DS710 interferents study showed no reactivity to common interferents present in sweat and urine.

**SI Figure 17.**
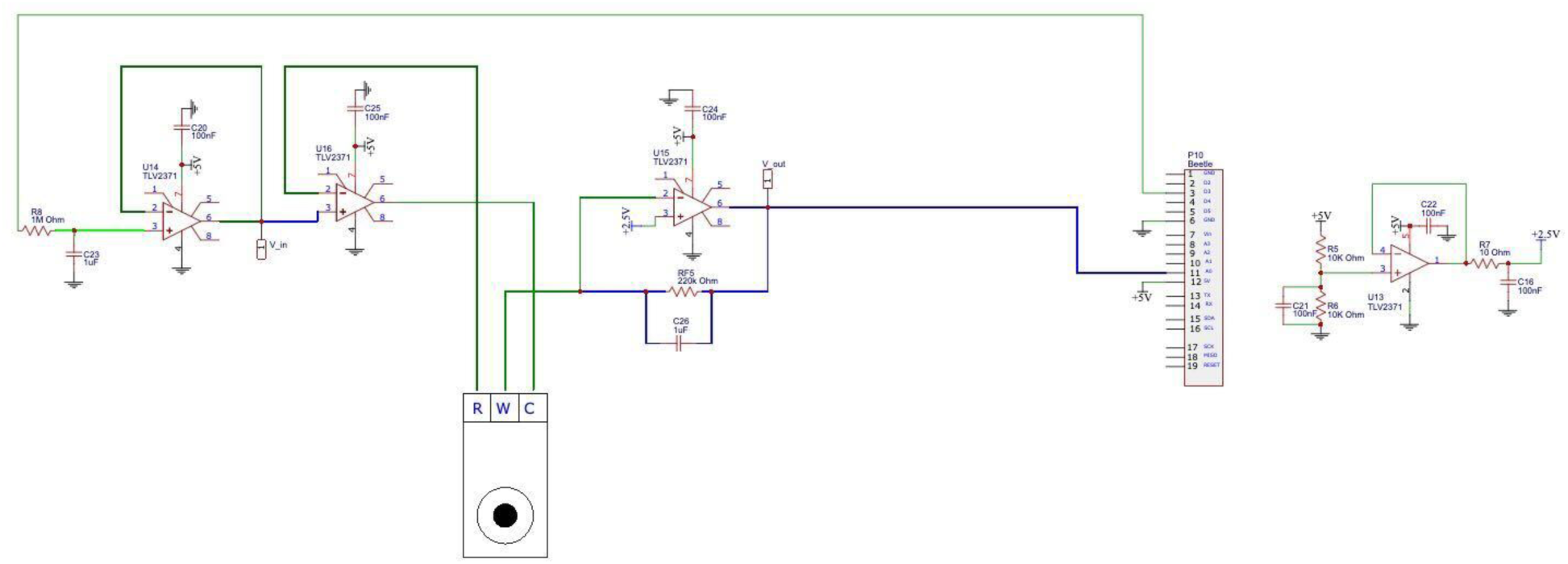
WearStat potentiostat circuitry which allows for the accurate measurement of a current response resulting from analyte addition to the electrochemical biosensor.

**SI Figure 18.**
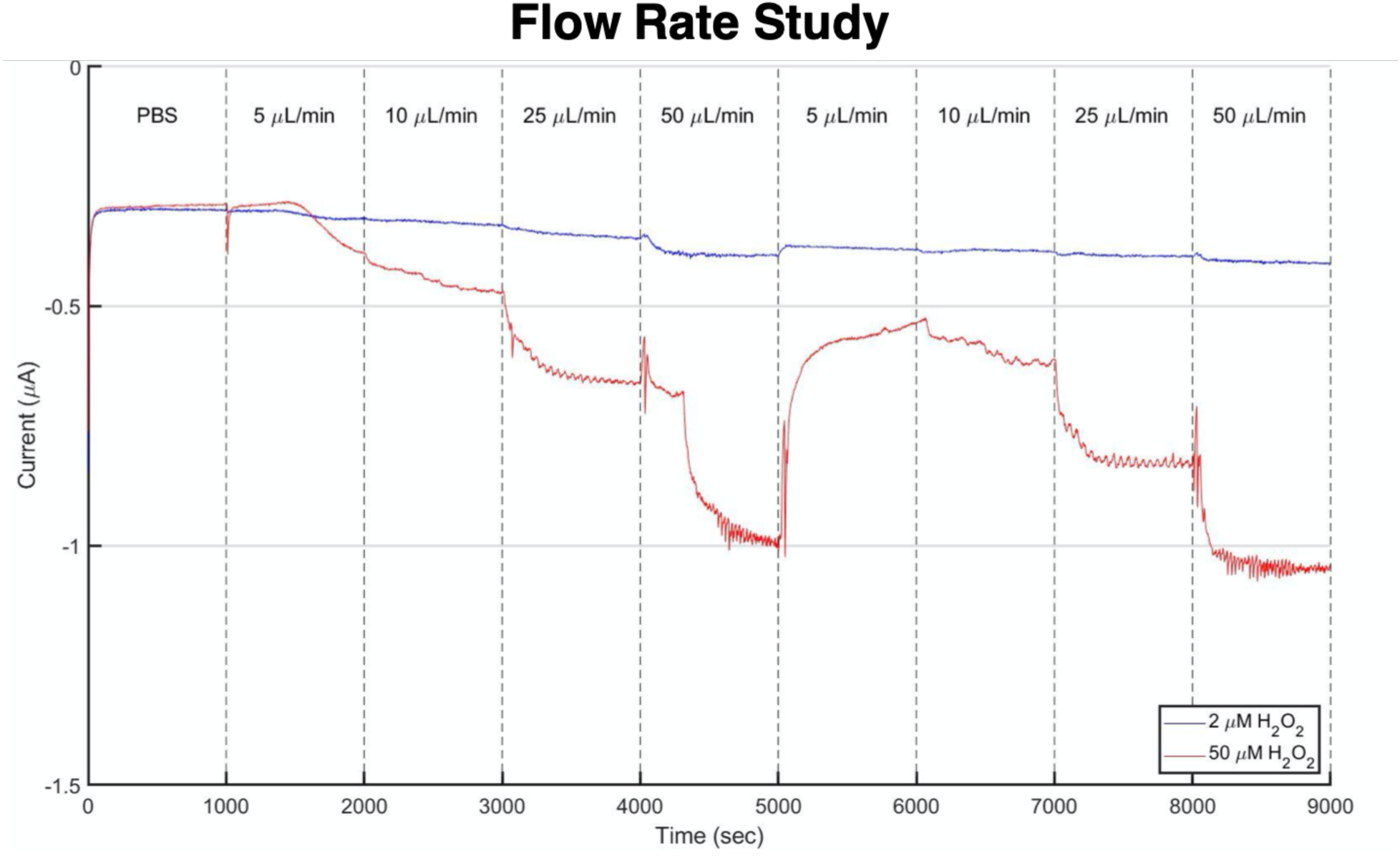
Analyte flow rate determines the current output. A syringe pump administered two H_2_O_2_ concentrations at four flow rates over two cycles onto a WearStat paper channel with a connected DS710 SPE.

**SI Figure 19.**
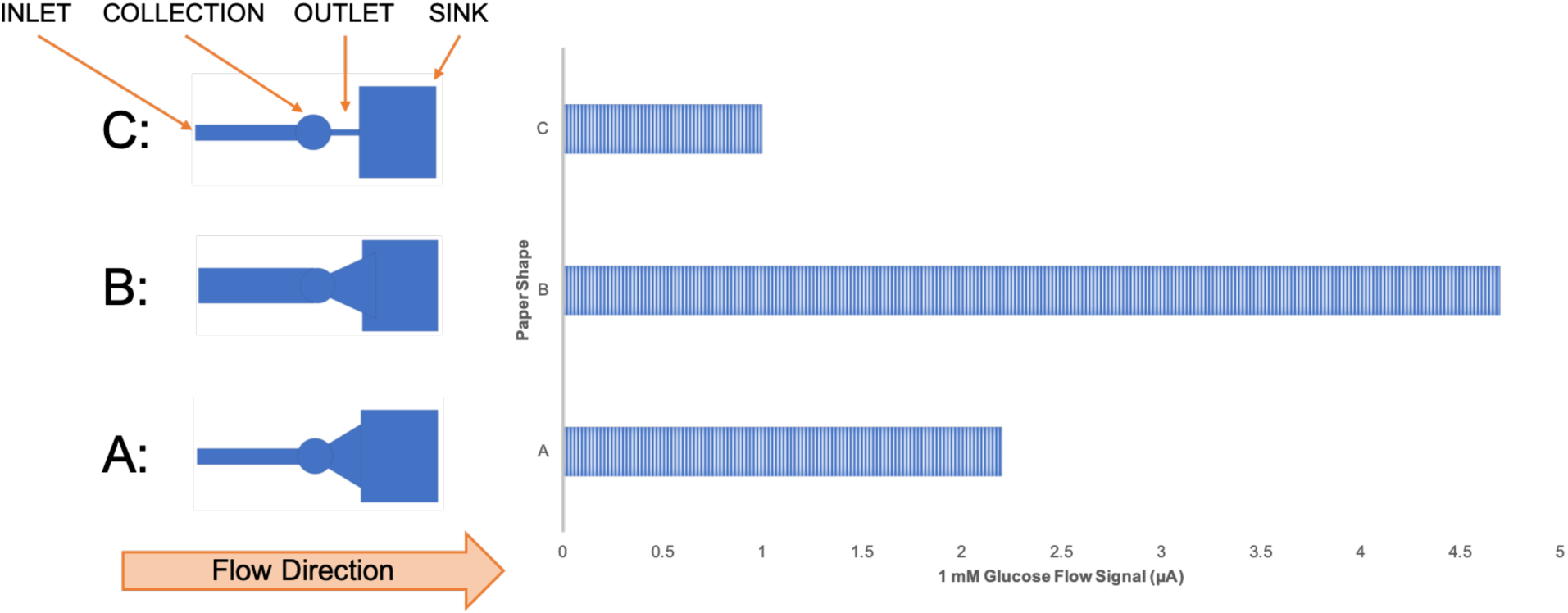
Paper channel geometries affect the flow rate and thus current output. A paper channel with a wide inlet and wide outlet proved to produce the largest current output. The collection and sink area remained constant through the three designs. Current was calculated as a delta in comparison to the baseline.

**SI Figure 20.**
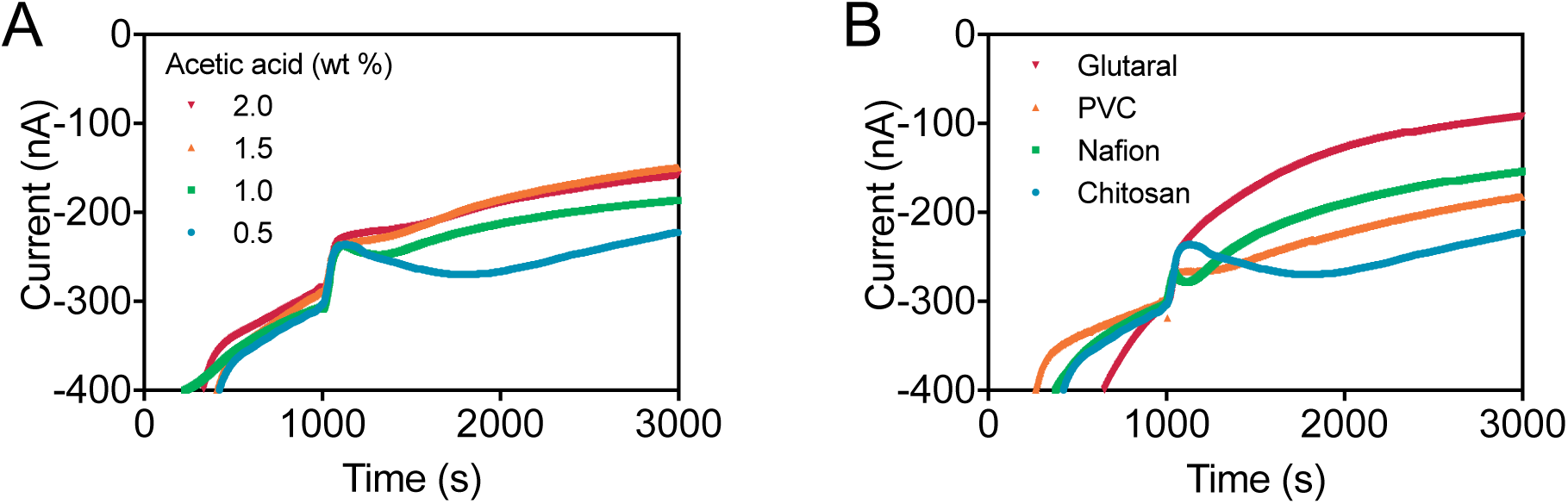
Chronoamperometric responses of the nicotine sensors with different immobilization methods. **(A)** Chronoamperometric responses with 1 wt% chitosan in four different weight percent acetic acid as immobilization matrix. **(B)** Effect of different immobilization methods. Four immobilization methods were used: a) 1 wt% chitosan in 0.5 wt% acetic acid with enzyme; b) Enzyme + 1% Nafion; c) 1 wt% chitosan in 0.5 wt% acetic acid in 10 mg/mL BSA with enzyme + 3 wt% PVC in tetrahydrofuran; d) 1 wt% chitosan in 0.5 wt% acetic acid in 10 mg/mL BSA with enzyme + 2 wt% glutaraldehyde. Currents were measured with 400 μM nicotine in PBS added to sensor prepared with different immobilization methods.

**SI Figure 21.**
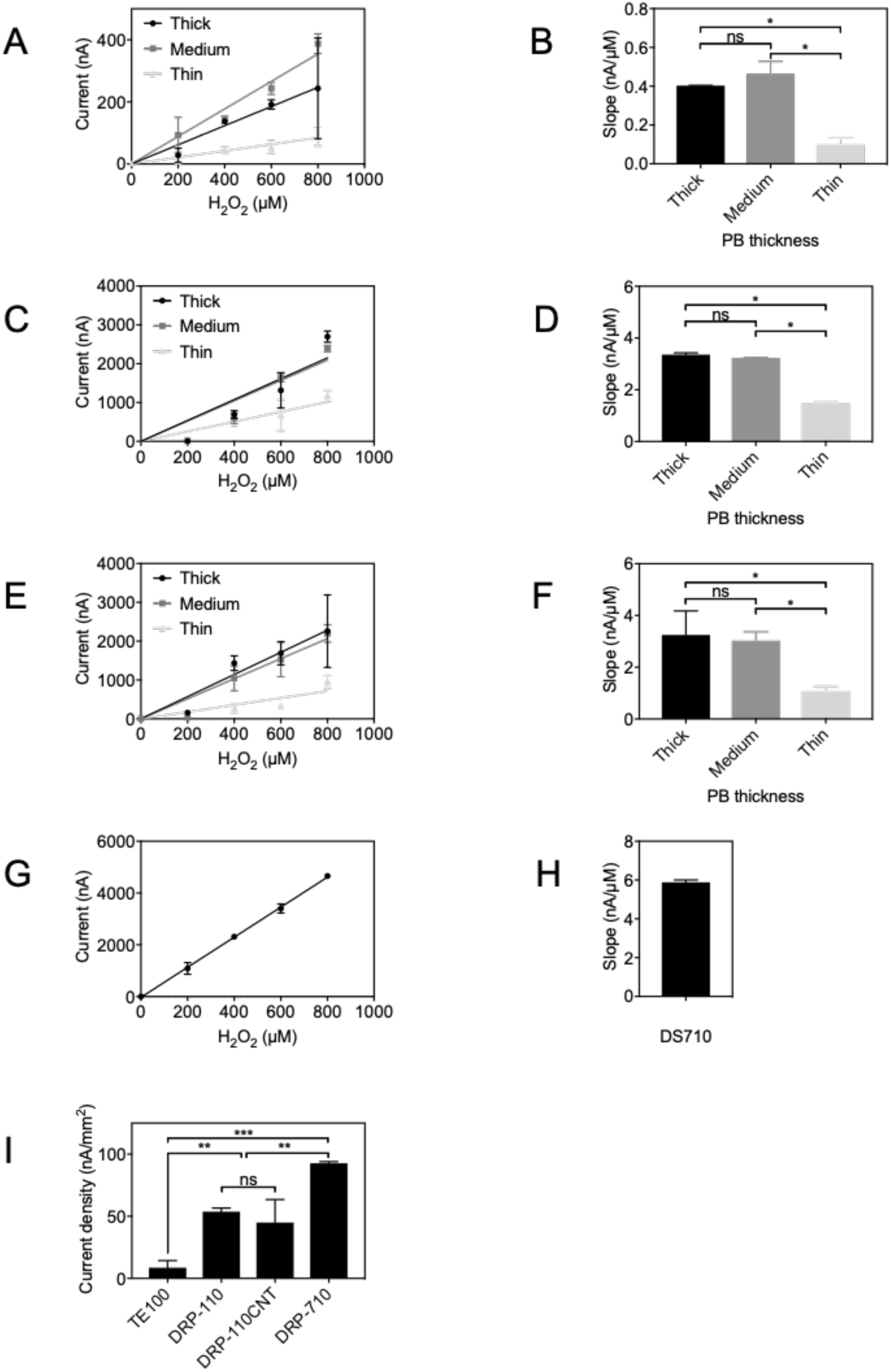
Screen-printed electrodes comparison. Dose-response curves for hydrogen peroxide detection using screen-printed electrodes modified with Prussian Blue: Prussian Blue was electrodeposited on **(A)** electrode TE100, **(C)** electrode DRP-110, **(E)** electrode DRP-110CNT with varying thickness; **(G)** DRP-710 electrodes pre-impregnated with Prussian Blue were used as received without further PB deposition. Hydrogen peroxide detection sensitivity of commercial screen-printed electrodes: **(B)** TE100, **(D)** DRP-110, **(F)** DRP-110CNT, and **(H)** DRP-710. **(I)** Current density of TE100 with medium PB, DRP-110 with thick PB, DRP-110CNT with thick PB, and DRP-710 at 800 μM hydrogen peroxide. *Indicates statistical difference (95% confidence) between samples.

**SI Figure 22.**
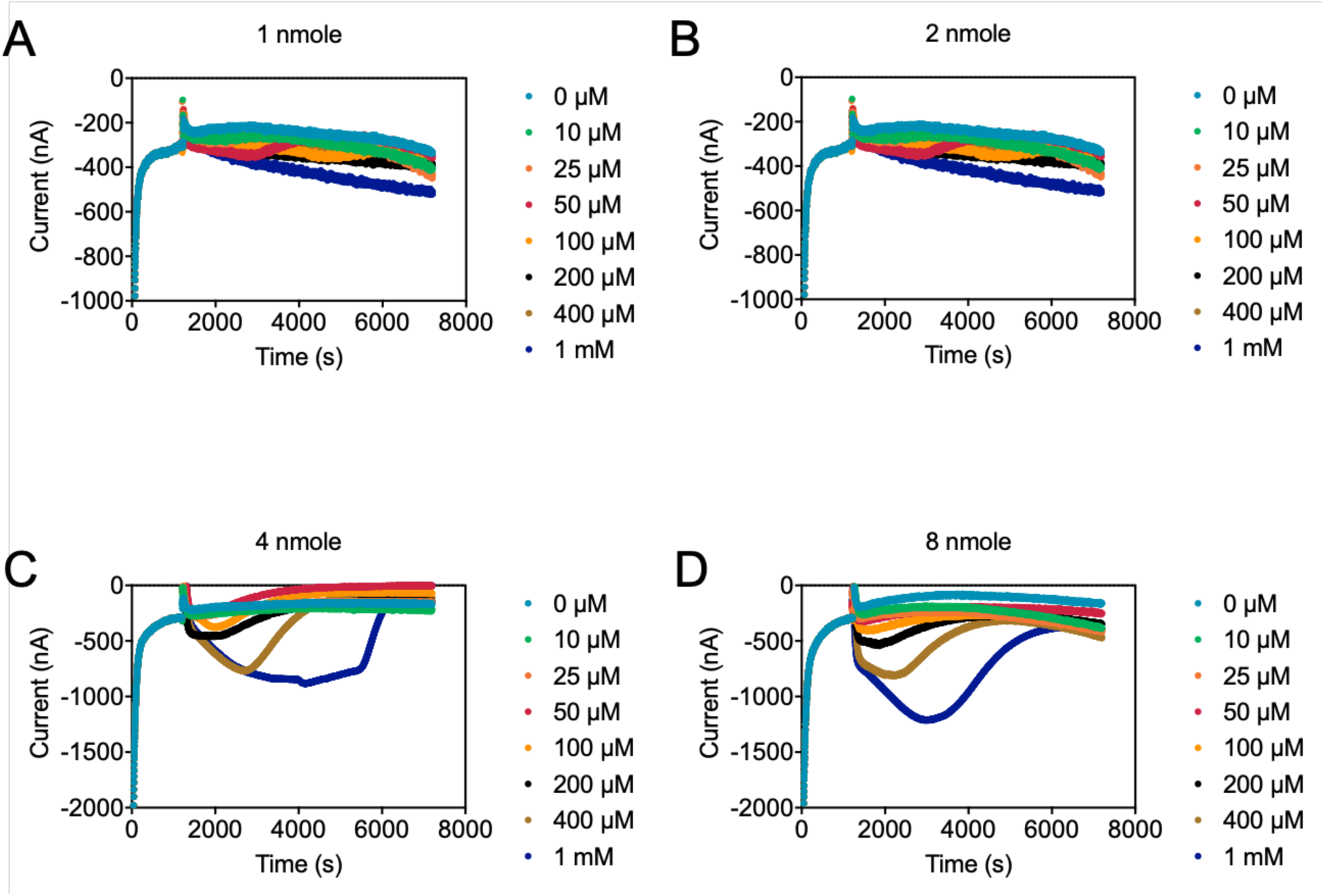
Chronoamperometric responses of the nicotine sensors with different enzyme amounts. **(A)** 1 nmol NicA2. **(B)** 2 nmol NicA2. **(C)** 4 nmol NicA2. **(D)** 8 nmol NicA2.

**SI Figure 23.**
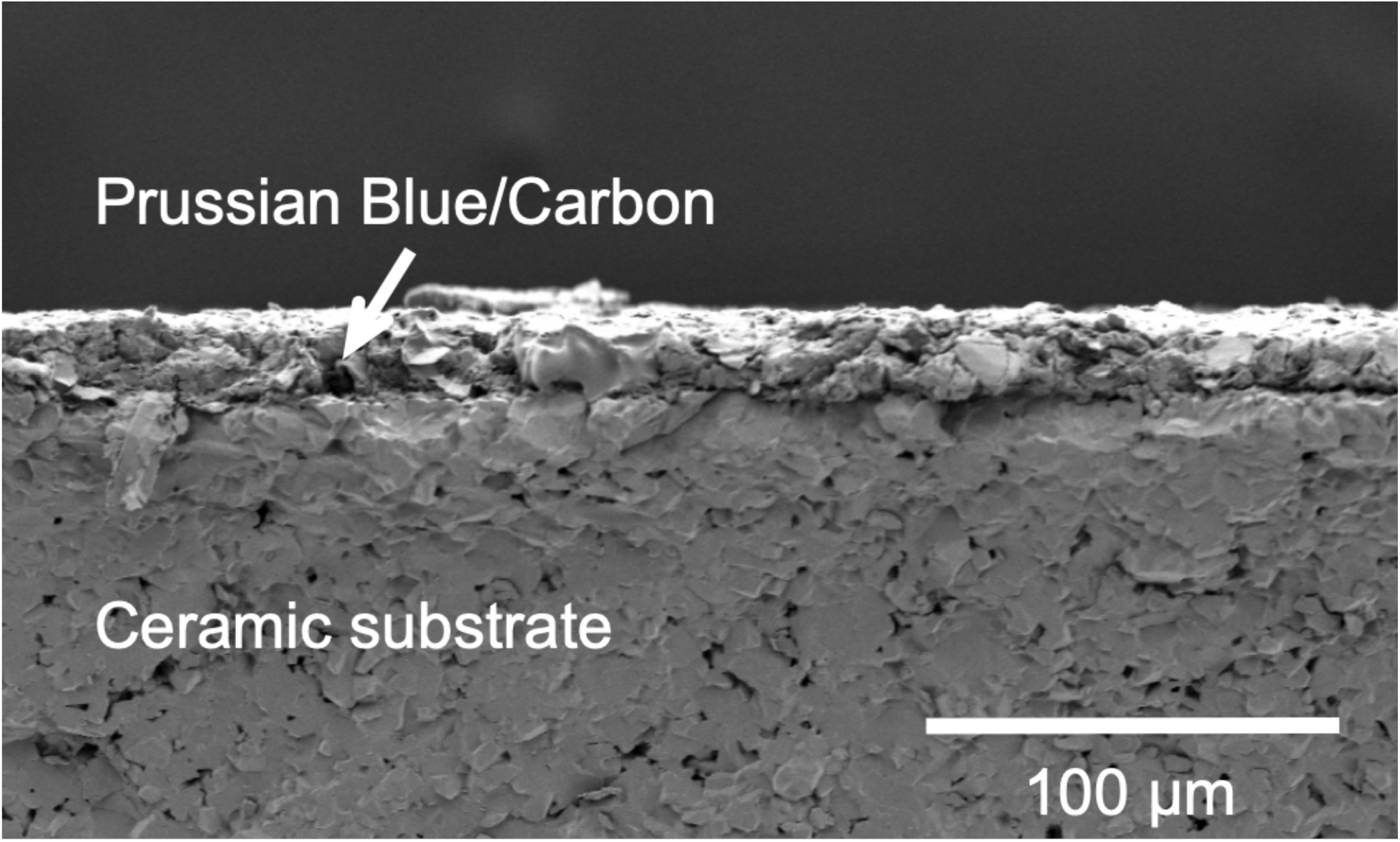
SEM image of DRP-710 cross section.

**SI Figure 24.**
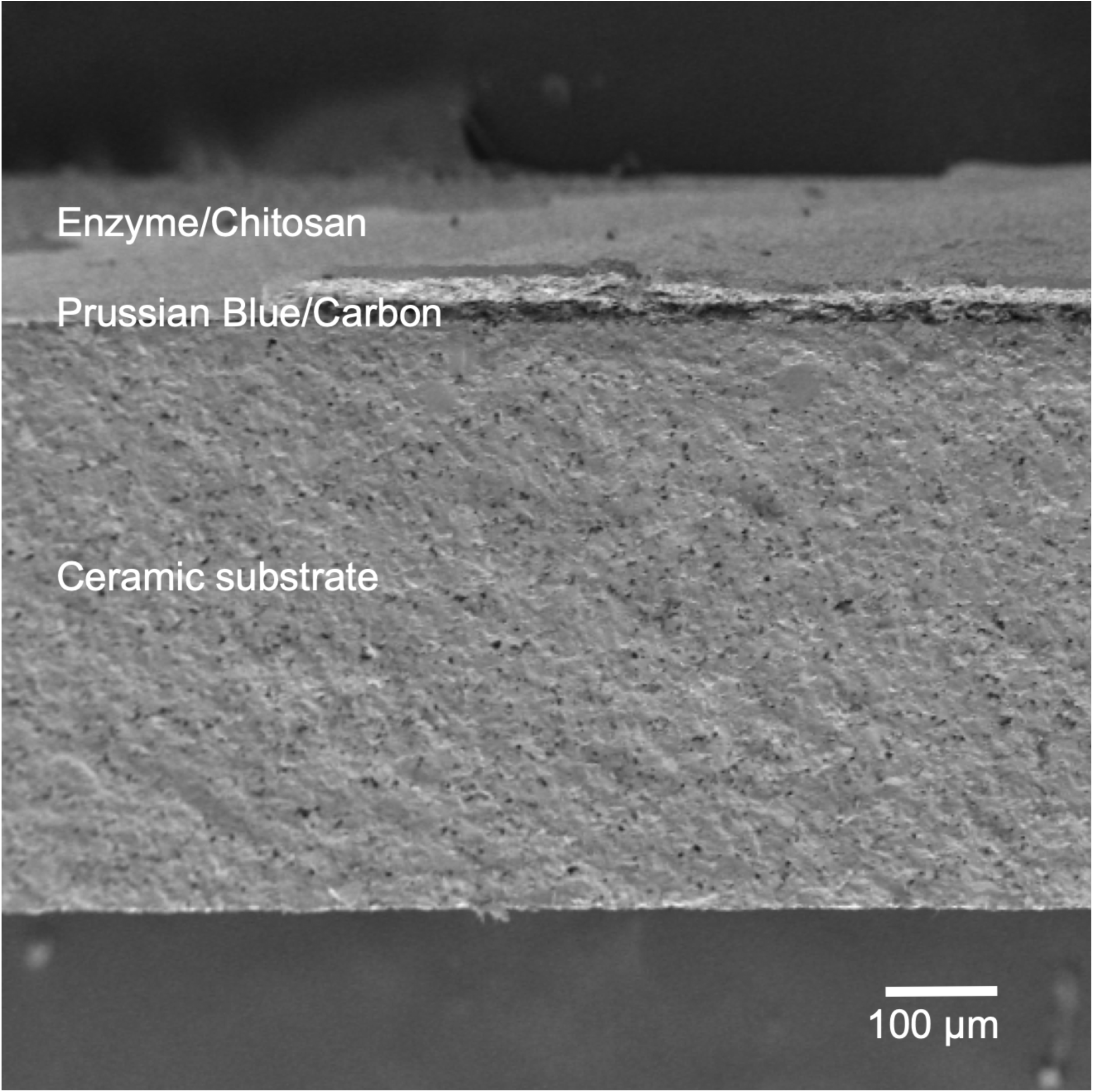
SEM image of nicotine sensor cross section.

**SI Figure 25.**
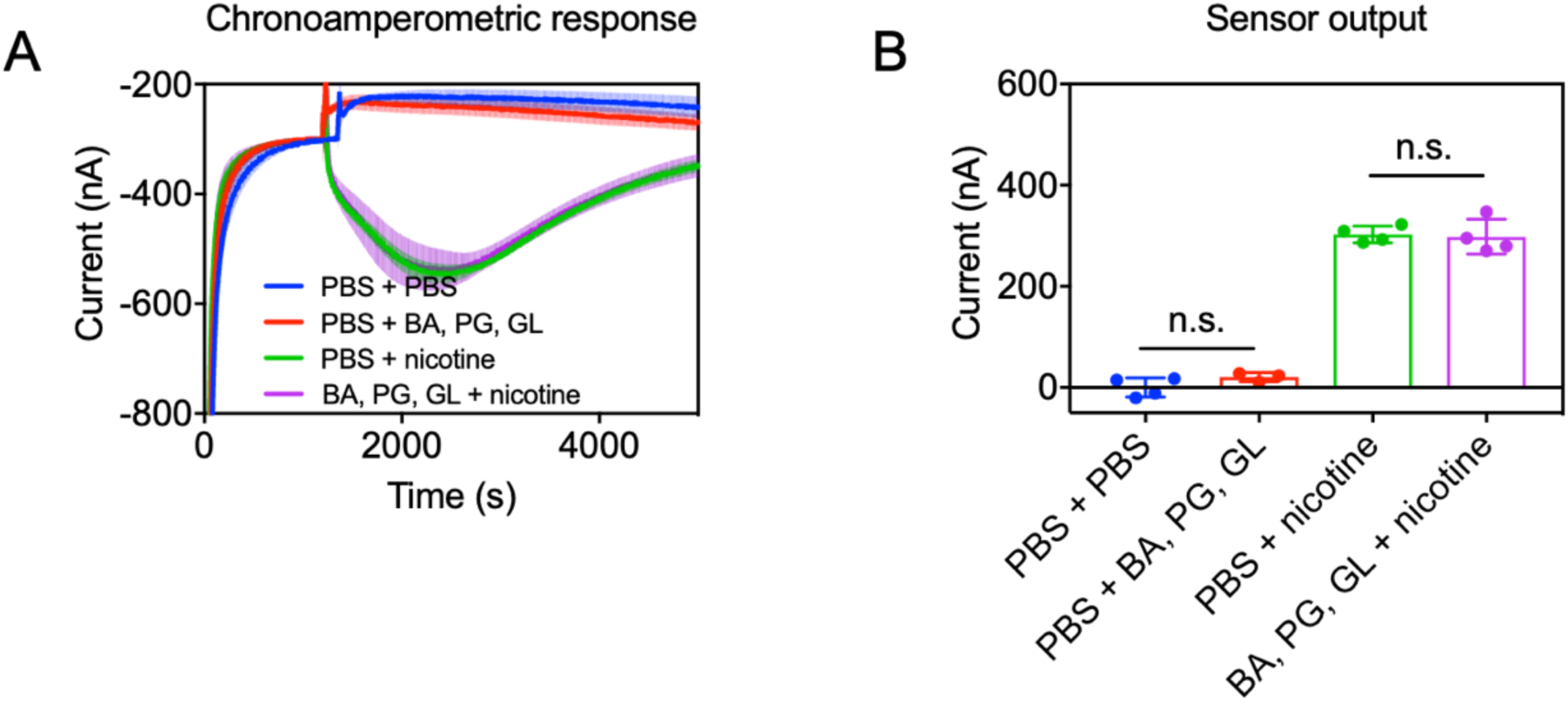
Selectivity of the sensor against common interferents in e-cigarette liquid. **(A)** Chronoamperometric response to common interferents BA, PG, GL: benzoic acid, propylene glycol, and glycerol. **(B)** Sensor outputs of the selectivity tests, as presented in A.

## Supplementary Tables

**SI Table 1.**
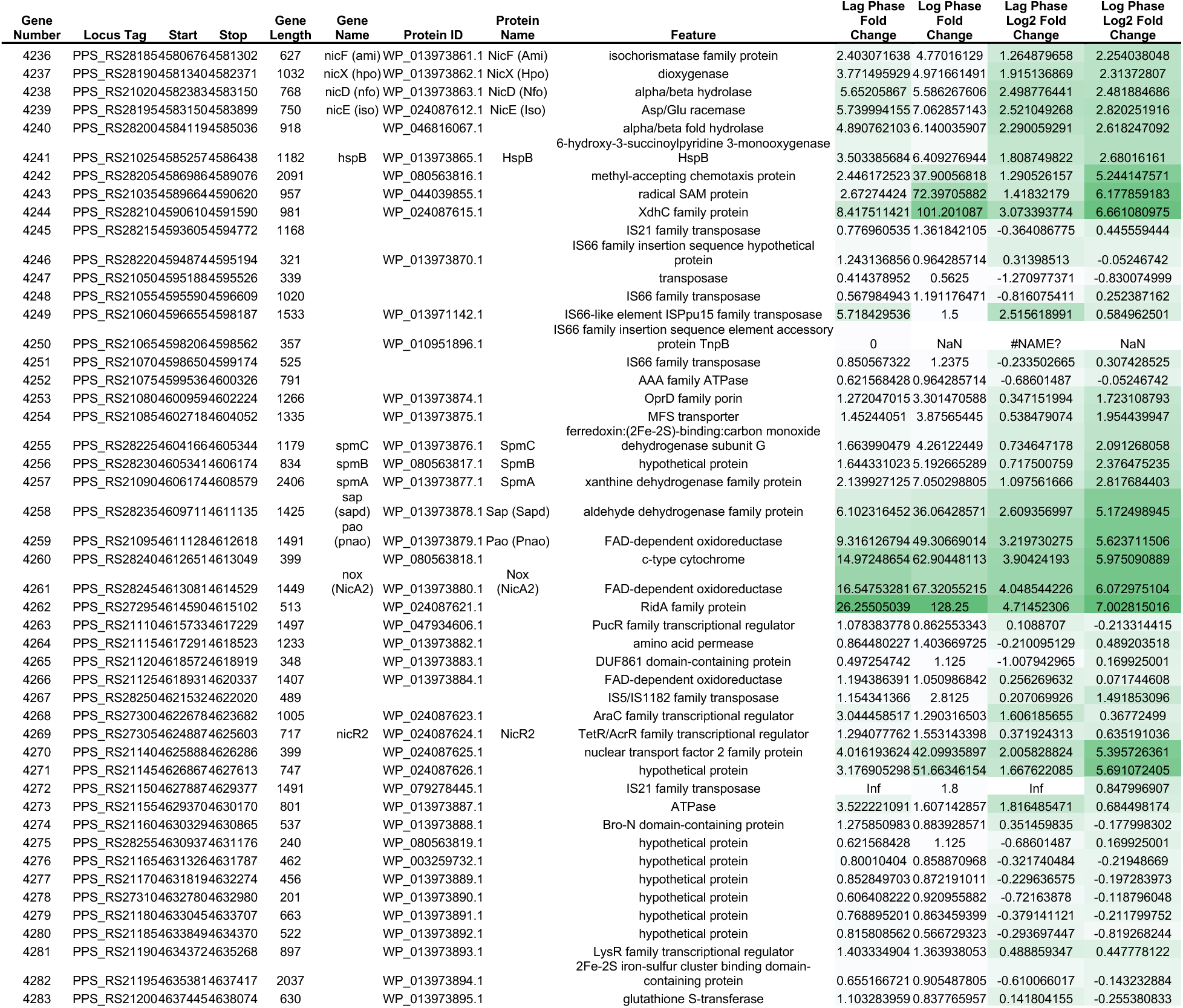
RNA-Seq quantitative differential expression of NRGI genes in the presence of nicotine compared to control during lag and log phases of growth.

**SI Table 2.** NicA2 is a more selective, yet slower redox enzyme compared to GO_X_ and LO_X_.

| Enzyme | $K_M$ | $k_{cat}$ | $k_{cat}/K_M$ | $1/k_{cat}$ |
| --- | --- | --- | --- | --- |
| NicA2 | 43.5 ± 4.7 nM | (6.64 ± 0.17) * 10<sup>-3</sup> s<sup>-1</sup> | 152.64 s<sup>-1</sup>mM<sup>-1</sup> | 150.6024 s |
| Glucose Oxidase (GO<sub>x</sub>) | 33-110 mM | 1118.81 s<sup>-1</sup> | 22.04 s<sup>-1</sup>mM<sup>-1</sup> | 0.000894 s |
| Lactate Oxidase (LO<sub>x</sub>) | 0.94 mM | 280 s<sup>-1</sup> | 297.87 s<sup>-1</sup>mM<sup>-1</sup> | 0.00357 s |

**SI Table 3.**
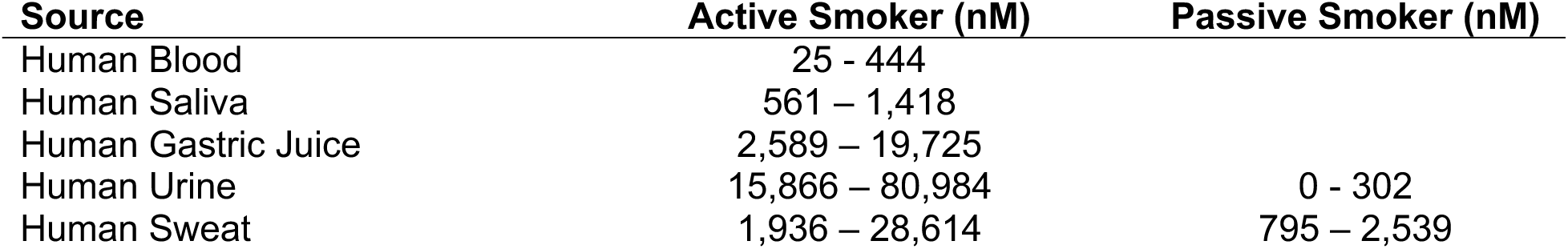
Nicotine levels in physiologically relevant human matrices. ^37,77,95–98^. The concentrations in red are the concentrations that can be detected by the enzyme-based nicotine biosensor.

| **Source** | **Active Smoker (nM)** | **Passive Smoker (nM)** |
| --- | --- | --- |
| Human Blood | 25 - 444 |  |
| Human Saliva | 561 – 1,418 |  |
| Human Gastric Juice | 2,589 – 19,725 |  |
| Human Urine | 15,866 – 80,984 | 0 - 302 |
| Human Sweat | 1,936 – 28,614 | 795 – 2,539 |

**SI Table 4.**
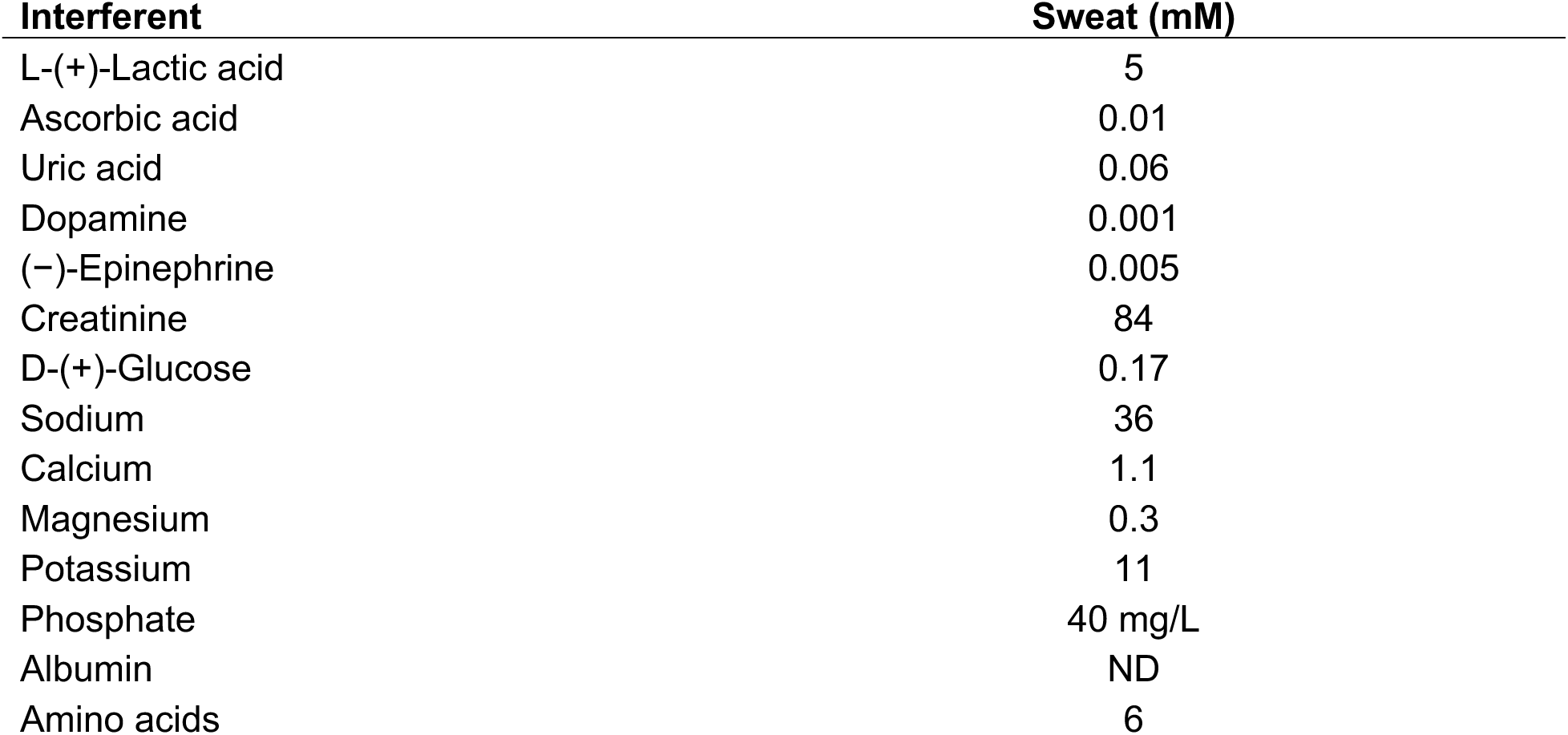
Interferents and their estimated concentration in sweat^8^.

| **Interferent** | **Sweat (mM)** |
| --- | --- |
| L-(+)-Lactic acid | 5 |
| Ascorbic acid | 0.01 |
| Uric acid | 0.06 |
| Dopamine | 0.001 |
| (-)-Epinephrine | 0.005 |
| Creatinine | 84 |
| D-(+)-Glucose | 0.17 |
| Sodium | 36 |
| Calcium | 1.1 |
| Magnesium | 0.3 |
| Potassium | 11 |
| Phosphate | 40 mg/L |
| Albumin | ND |
| Amino acids | 6 |

**SI Table 5.**
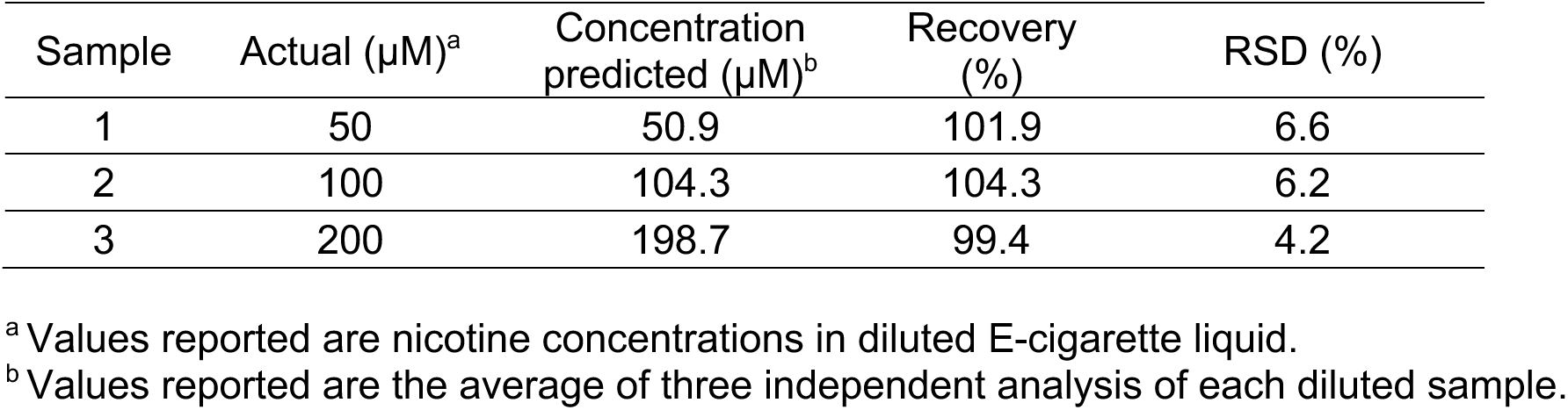
Nicotine recovery experiments in diluted E-cigarette liquid.

| Sample | Actual ($\mu\text{M}$)<sup>a</sup> | Concentration predicted ($\mu\text{M}$)<sup>b</sup> | Recovery (%) | RSD (%) |
| --- | --- | --- | --- | --- |
| 1 | 50 | 50.9 | 101.9 | 6.6 |
| 2 | 100 | 104.3 | 104.3 | 6.2 |
| 3 | 200 | 198.7 | 99.4 | 4.2 |

<sup>a</sup> Values reported are nicotine concentrations in diluted E-cigarette liquid.
<sup>b</sup> Values reported are the average of three independent analysis of each diluted sample.

**SI Table 6.**
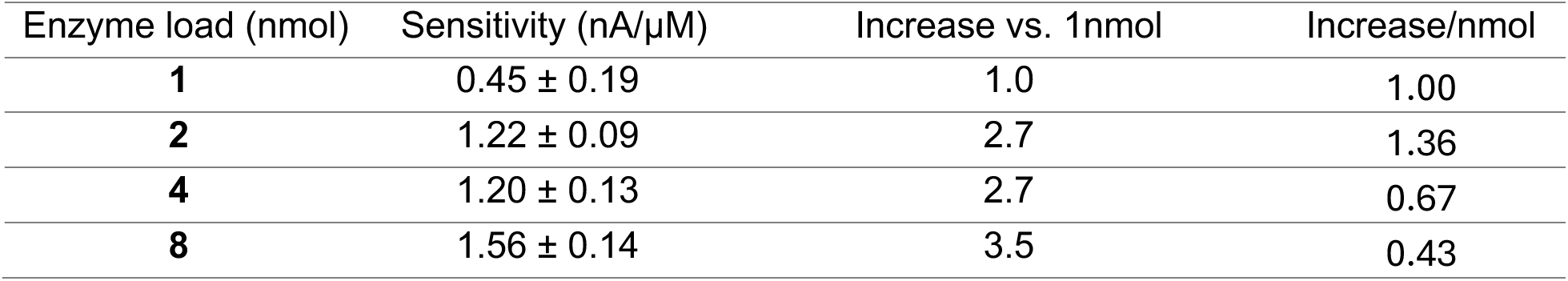
Dose response in the range of 0 to 200 μM nicotine with the presence of increasing enzyme load.

| Enzyme load (nmol) | Sensitivity (nA/μM) | Increase vs. 1nmol | Increase/nmol |
| --- | --- | --- | --- |
| **1** | 0.45 ± 0.19 | 1.0 | 1.00 |
| **2** | 1.22 ± 0.09 | 2.7 | 1.36 |
| **4** | 1.20 ± 0.13 | 2.7 | 0.67 |
| **8** | 1.56 ± 0.14 | 3.5 | 0.43 |

**SI Table 7.** Cross section thickness of nicotine sensor constituent layers, measured with SEM.

|  | **Thickness ($\mu\text{M}$)** | **Standard Deviation ($\mu\text{M}$)** |
| --- | --- | --- |
| Enzyme layer | 95.0 | 1.2 |
| Prussian Blue/Carbon layer | 22.5 | 0.8 |
| Ceramic substrate | 514.8 | 7.0 |

**SI Table 8.**
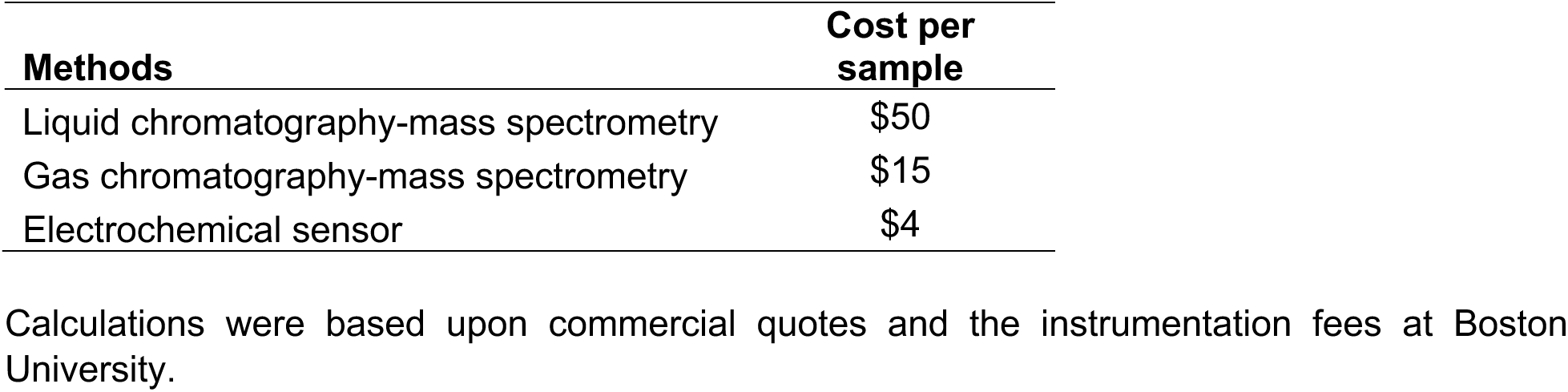
Cost of electrochemical sensor compared to traditional analytical methods.

| **Methods** | **Cost per sample** |
| --- | --- |
| Liquid chromatography-mass spectrometry | \$50 |
| Gas chromatography-mass spectrometry | \$15 |
| Electrochemical sensor | \$4 |

Calculations were based upon commercial quotes and the instrumentation fees at Boston University.

## Author Acknowledgements

The authors would like to acknowledge the BU Biomedical Engineering Core Facilities (BECF), the BU Chemical Instrumentation Center (CIC), the BU Optoelectronic Processing Facility (OPF), and the BU Center for Nanoscale Systems (CNS). The authors would also like to acknowledge Stephen Whelan, Maryam Salari, Andy Fan, Takumi Hawes, and Nicolas Shu for their contributions. Z.Z., A.A., H.Z. U.K., R.C., and J.E.G received support from DOD/ONR (Awards N000142112433, N000142212689); A.A., H.Z., and J.E.G. received support from DASA (Contract No. DSTL0000016713); U.K., R.C., J.E.G. received support from NSF (Award 2126992), MC received support from a BU BME distinguished fellowship, MC and UK received support from a BUnano Terrier Tank award.

## Author Contributions

Based on https://www.niso.org/publications/z39104-2022-credit

Conceptualization: U.K., M.C., C.K., K.N.A., M.W.G., J.E.G.

Data curation: U.K., M.C., R.C.

Formal analysis: U.K., M.C., K.H., J.E.G.

Funding acquisition: U.K., C.K., K.N.A., M.W.G., J.E.G.

Investigation: U.K., M.C., R.C., P.S., A.G., S.D, N.L.G.D, K.S., K.H., H.Z.

Methodology: U.K., M.C., J.E.G.

Project administration: C.K., K.N.A., M.W.G., J.E.G.

Resources: K.H., M.A.T, A.M.D, H.Z.

Software: U.K., M.C., A.A., J.E.G.

Supervision: C.K., K.N.A., M.W.G., J.E.G.

Visualization: U.K., M.C., J.E.G.

Writing: U.K., M.C., Z.Z., A.A., K.N.A., M.W.G., J.E.G.

Reviewing and Editing: U.K., M.C., K.N.A., M.W.G., J.E.G.

